# Unsupervised manifold learning using low-distortion alignment of tangent spaces

**DOI:** 10.1101/2024.10.31.621292

**Authors:** Dhruv Kohli, Johannes S. Nieuwenhuis, Katja Zegwaard, Alexander Cloninger, Gal Mishne, Devika Narain

## Abstract

With the ubiquity of high-dimensional datasets in various biological fields, identifying low-dimensional topological manifolds within such datasets may reveal principles connecting latent variables to measurable instances in the world. The reliable discovery of such manifold structure in high-dimensional datasets can prove challenging, however, largely due to the introduction of distortion by leading manifold learning methods. The problem is further exacerbated by the lack of consensus on how to evaluate the quality of the recovered manifolds. Here, we present a novel measure of distortion to evaluate low-dimensional representations obtained using different techniques. We additionally develop a novel bottom-up manifold learning technique called Riemannian Alignment of Tangent Spaces (RATS) that aims to recover low-distortion embeddings of data, including the ability to embed closed manifolds into their intrinsic dimension using a unique tearing process. Compared to previous methods, we show that RATS provides low-distortion embeddings that excel in the visualization and deciphering of latent variables across a range of idealized, biological, and surrogate datasets that mimic real-world data.

**One-sentence summary:** We introduce a novel dimensionality reduction technique that generates low-dimensional embeddings while preserving the global structure within the data for a variety of biological and non-biological datasets.

## Introduction

The manifold hypothesis is prevalent in many biological and non-biological fields, where complex, multifaceted datasets are thought to be well-represented by latent lower-dimensional manifolds despite being observed in high-dimensional feature spaces. Such manifold-like structures have emerged in several empirical applications within computational domains such as computer vision^1^, language and representation learning^2–4^, and in biological domains such as gene expression^5–8^, oncology^9^, botany^10^, and neuroscience^11–16^. These structures are often revealed using manifold learning techniques, which aim to learn a lower-dimensional representation of data, ideally capturing the *intrinsic dimension* of the dataset. The intrinsic dimension refers to the minimum number of variables needed to effectively describe the data^17,18^, *e*.*g*., the intrinsic dimension of a sphere is two, even though it is embedded in three dimensions.

Recent research in biological and non-biological fields^19–22^ has demonstrated valuable insights gained through the application of unsupervised manifold learning techniques and their derivatives, such as t-distributed Stochastic Neighbor Embedding^23^ (t-SNE), Uniform Manifold Approximation and Projection^24^ (UMAP), Isometric feature mapping^25^ (ISOMAP), Local Tangent Space Alignment^26^ (LTSA), Low Distortion Local Eigenmaps^27^ (LDLE), Hessian Locally Linear Embedding (hLLE)^28^ Maximum Variance Unfolding (MVU)^29^, and Local Orthogonality Preserving Alignment (LOPA)^30^, among others. Recent work has also adopted supervised methods for manifold learning^31^. These techniques differ in their assumptions regarding preserved geometric properties and in their choice of trade-offs while computing low-dimensional embeddings. These methods frequently incur significant distortion of distances along the original data manifold when mapping into the embedding space. Such distortion often arises from the uneven expansion or contraction of local spaces around embedded data points^32^ (Figure 1a). This is problematic for example, when we expect the latent variables of the manifold to have physical meaning, such as space or time. In cases where the measured data does not fully represent such physical quantities, we may be unable to disambiguate such a scenario from methods that inadvertently introduce distortions, resulting in a skewed interpretation of relationships with external variables. This can have undesirable effects, such as reduced reliability in decoding how biological variables represent phenotypes and behaviors. Consequently, erroneous conclusions may be drawn due to such algorithmic artifacts. In this work, we seek to mitigate this problem in two ways. First, we present a novel measure to quantify distortion in low-dimensional embeddings across different manifold learning techniques. Second, we develop a new bottom-up manifold learning method called Riemannian Alignment of Tangent Spaces (RATS) for the unsupervised discovery of low-distortion embeddings of complex datasets.

**Figure 1.**
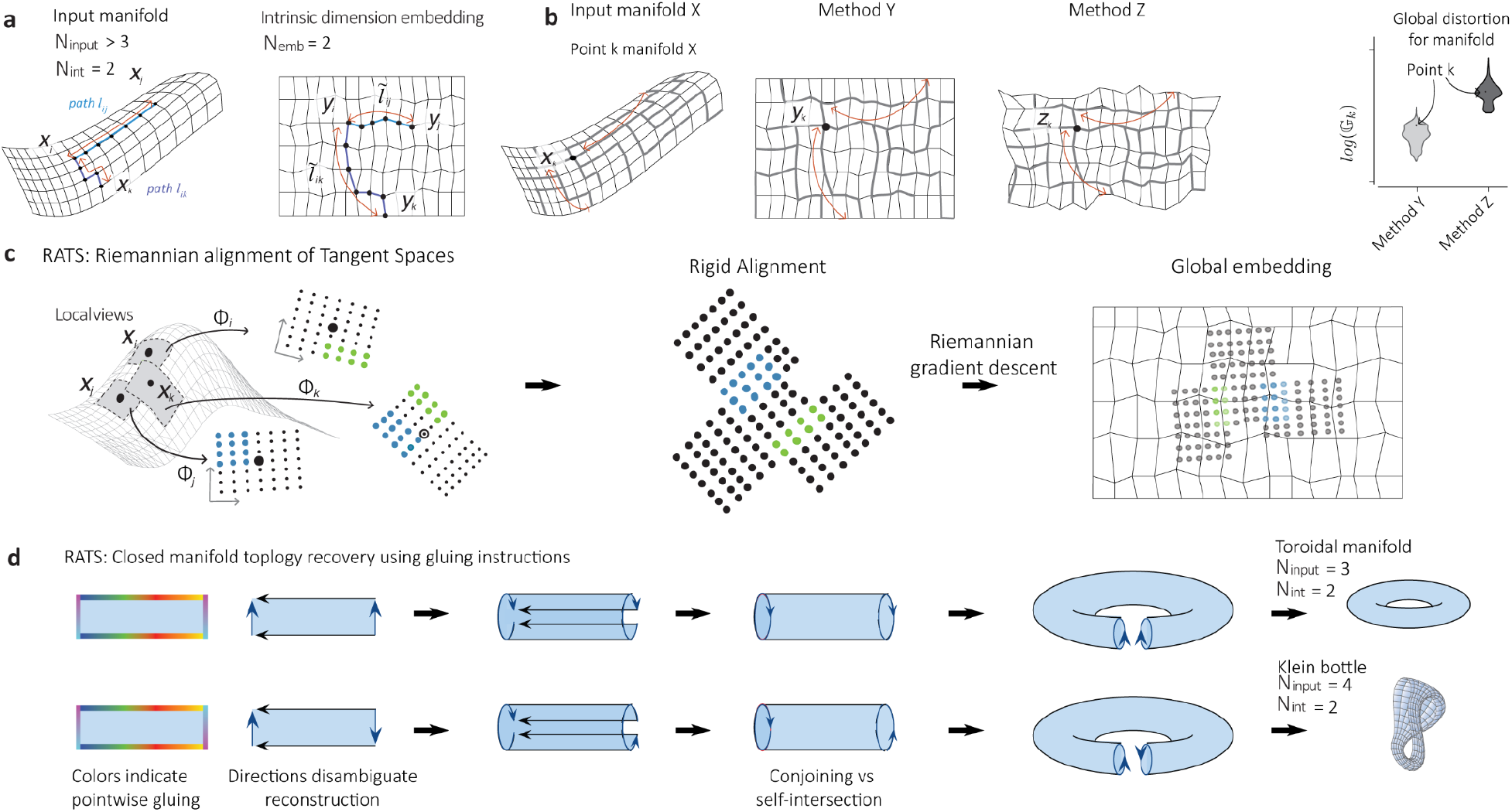
Global distortion metric and RATS with topology recovery. a) Embedding data points from an ambient dimension (N_input_) into a low-dimensional embedding space (N_emb_), which corresponds to the intrinsic dimension of the data (N_int_). b) Embeddings can produce distortion due to contraction and expansion of relative distances along geodesic paths. Here, x_k_ represents data point k on the original manifold, and y_k_ and z_k_ represent its counterparts in two different embeddings. For each datapoint, we calculate its geodesic distances to all other points in the original and the embedding space, and identify the paths with the worst contraction (path is shorter in the embedding) and expansion (path is longer in the embedding) compared to the original space. The product of these defines the global distortion at a point. Calculating the distribution of global distortion for all points in the datasets, allows us to quantitatively evaluate which embedding better preserves path lengths from the original space. c) RATS computes low-distortion global embeddings in two phases. First low-dimensional low-distortion local views of the dataset are constructed. Second these are aligned, based on their overlaps (blue and green points), using Riemannian gradient descent to obtain a global embedding. d) RATS is capable of tearing closed/non-orientable manifolds in order to embed them into their intrinsic dimensions. When RATS tears the input manifold, the method provides gluing instructions at the output by coloring the points along the tear so that the same colored points are to be stitched together. The arrows constituting the gluing diagram are recovered by tracking the variation of the colors. The schematic shows the recovery of the topology of a torus versus a Klein bottle through different gluing diagrams on the same 2D embedding.

### A novel measure of distortion

While low dimensional embeddings are ubiquitous, an open challenge is quantifying their quality. Some evaluation approaches utilize downstream tasks^19^, for example evaluating clustering accuracy in the embedding space^19,33,34^. Other measures evaluate the preservation of local information, *e*.*g*, calculating the percentage of k-nearest neighbors in the original high-dimensional data that are preserved in the embedding^35,36^. However, this is insufficient, since even if the ranking or composition of nearest neighbors is preserved, the distances between points can still be distorted. Local distortion ^32,37^ measures the contraction and expansion of Euclidean distances of local neighborhoods in the embedding with respect to distances of those neighborhoods in the ambient space. However, a quantitative analysis of the *global* distortion, i.e. preservation of the global geometry of the data in the embedding space, is lacking in the literature.

Here, we present a natural way to quantify the global distortion introduced when embedding data into a low-dimensional space based on the disparity between the geodesics (shortest paths between pairs of points) in the data 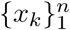 and the corresponding geodesics in the low-dimensional embedding, 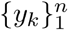 (Figure 1a,b). In general, the geodesic distances in the embedding differ from their counterparts in the data due to warping induced when embedding data into lower dimensions. We consider that any distortion between these distances arises from expansions and contractions of geodesic lengths rather than from isometric scaling. To capture such distortion, we present a novel measure of *Global distortion*, 𝔾_*k*_, defined at the *k*th point, as the product of the worst expansion and the worst contraction of the shortest paths originating from *x*_*k*_ (Figure 1b). This measure is invariant to isometric transformations and scaling of the data, *i*.*e*. rotating the dataset or scaling all the points by a constant will result in global distortion of 1 for all points. Considering the global distortion of all points in the data generates a distribution of distortions (Figure 1b), enabling a more comprehensive evaluation of embeddings. It should be noted that this method relies on accurately calculating geodesic lengths, which is challenging in the presence of significant noise. However, we recommend mitigating such sensitivity to noise by applying denoising algorithms ^38^ to the input data.

### Manifold learning through Riemannian Alignment of Tangent Spaces (RATS)

To target the challenge of generating embeddings with low distortion, we present a new bottom-up manifold learning technique called *Riemannian Alignment of Tangent Spaces* (RATS, Figure 1c-d). Unlike classical manifold learning approaches, which typically solve for a single global embedding through a single optimization or eigenvalue problem, RATS is a bottom-up approach that first generates low-distortion local views of the data. Construction of these views can accommodate different similarity measures (*e*.*g*., Euclidean, cosine) to obtain a local parametrization of each point. RATS then learns a rigid alignment (rotations and translations) between these views, based on overlapping points between pairs of views, using Riemannian Gradient Descent^39^. This yields a single global embedding of the data. A key insight here is that for bottom-up manifold learning, ensuring low local distortion in generating local views and aligning them optimally, *i*.*e*. with small error, will result in minimal accrual of distortion in the global embedding. Thus, while RATS does not optimize a cost function based on Global distortion 𝔾_*k*_ to obtain an embedding, the design of its construction process will result in a global embedding with low global distortion.

Furthermore, RATS offers an optional feature for embedding manifolds that is rarely present in existing manifold learning methods. It can tear closed manifolds in order to embed these in their intrinsic dimension^17^ and provide “gluing” instructions to piece the torn manifold into its original closed topology (Figure 1d). This tearing strategy is especially useful for decoding external or latent variables from the data because it lets users read off parameters simply and directly from the manifold’s intrinsic coordinates, rather than inferred from extra dimensions introduced by curvature. For example, the intrinsic dimension of a circle is one, being parametrized by a single variable, the radial angle. Most manifold learning techniques, however, require two parameters to recover an underlying circle from data (*i*.*e*., they recover sine and cosine of the angle rather than the angle itself). Similarly, one’s location on Earth can largely be described by two coordinates, latitude and longitude, and an atlas or map of the Earth is a representation that tears the manifold surface to embed it in two dimensions. For many manifold learning techniques, the attempt to embed a sphere into two dimensions will result in a collapsed manifold, where for instance, the north and the south hemispheres lie on top of each other in 2D (Supplementary figure 1a-b). In such a case, we obtain an undesirable many-to-one mapping (*i*.*e*., both the north pole and south pole are mapped to the same point), and lose information that enables the recovery of the original points in data space since distances are distorted (*e*.*g*., Japan and Australia are artificially close in Supplementary figure 1a-b), thereby hindering our ability to interpret and decode such relationships. By enabling the option to tear and embed the manifolds in their intrinsic dimension, RATS preserves their intrinsic structure, preventing undesired overlap and enabling straightforward decoding of important parameters directly from the lower-dimensional manifold.

In addition to gluing instructions that enable tearing, RATS provides disambiguating instructions on how to stitch or reconstruct a torn manifold, so that its topology can be recovered (arrows in Figure 1d). For instance, a low-dimensional rectangular embedding may be folded into more than one plausible topological object, e.g., a hyperplane, a torus, or a Klein bottle. RATS can disambiguate these cases by providing a gluing diagram that can lead to the recovery of these two later distinct topologies, or in the case of the former, indicate that the manifold has not been torn. The gluing diagram directly represents the underlying topology, without needing to resort to alternative topological data analysis tools for validation. Moreover, we introduce a feature to optimize the tearing of manifolds using a cut-and-paste operation that can be applied to the RATS embedding (Supplementary Figure 2a-b, Methods).

Previous manifold learning techniques have aimed to address embedding data in low dimensions to mitigate distortions. However, many of these techniques cannot scale to large datasets easily due to computational constraints or do not handle noisy data well. For instance, ISOMAP seeks to preserve geodesic distances between points when embedding from high to low dimensions. hLLE^28^ and MLLE^40^, which are extensions of LLE^41^, can recover local isometry in many cases where methods such as ISOMAP fail, while MVU^29^, in addition to other related methods^42–44^, maximizes the spread of points and has theoretical convergence guarantees. Recent work includes LOPA^41^, which adapts LTSA^26^, to preserve both local distances and angles. However, treatment of noise, sparse data, and computational efficiency limit the utility and generalizability of these methods.

## Results

The true intrinsic dimension of manifold objects is often challenging to ascertain in many empirical datasets^17,18^, which are often plagued by inconsistencies, noise, and discontinuities, among other irregularities. An ideal low-distortion manifold learning technique would be able to perform well in a wide variety of circumstances, *i*.*e*., when the ground-truth topology and intrinsic dimensions are unknown, or where the presence of noise shrouds our understanding of the true topology of low-dimensional objects. We, therefore, approached the problem in three steps. First, we used ten idealized datasets, where the generative properties of the manifold were known, to evaluate RATS alongside other methods using the *Global distortion* measure 𝔾_*k*_ ^1^. Second, we used real-world datasets(N = 8) from six studies that investigated different behaviors in various brain structures. We examined how RATS and other manifold learning methods performed when the underlying topologies and sizes of latent representations of the dataset are unknown. We were able to evaluate 𝔾_*k*_ for a subset of these. Third, we used several deep neural network datasets to mimic naturalistic datasets where the underlying topology and task relevance could be determined but which also contained realistic features, such as noise, inhomogeneities, missing data, and sparsity. Using 𝔾_*k*_, and decoding of external variables where relevant, we were able to evaluate the performance of RATS and other methods on idealized, biological, and amalgamated datasets. Overall, using a quantifiable measure of distortion and methods that achieve low distortion does not only result in higher-quality embeddings for visualizations but may additionally reveal principles by which such topologies represent variables in the external world.

### Evaluating RATS on idealized datasets

To evaluate the benefit of a low-distortion method like RATS in uncovering latent structure, we first consider idealized datasets where ground truth topologies are established (Figure 2a-g). We compute 2D (*N*_emb_ = 2) embeddings of manifolds with varying dimensionality (barbell, *N*_int_ *≤ 2, N*_input_ *= 2*, Figure 2a), manifolds with missing data with distinct local geometries (square with cutouts, *N*_input_ = 2, Figure 2b), nonlinear manifolds with missing data (Swiss roll with a cutout, *N*_input_ = 3, Figure 2c) and with noise (Swiss roll with noise, *N*_input_ = 3, Figure 2d). For each dataset, we calculate the distortion in embeddings produced by various top-down techniques, *viz*., t-SNE^23^, UMAP^24^, ISOMAP^25^ and various bottom-up methods, viz., LTSA^26^, MVU^2945^, IMVU^46^, hLLE^28^, mLLE^40^, LDLE^27^, and RATS. After optimizing hyperparameters for each technique (Table 1, Supplementary figure 3a-e), a nonparametric analysis of variance (Kruskal-Wallis) reveals a significant main effect of global distortion across techniques for each of these datasets (Figure 2a-d, Supplementary table 1). RATS consistently exhibits the lowest global distortion in each of these idealized datasets. We demonstrate that for these idealized datasets, RATS-generated embeddings excel in preserving local geometric features, *i*.*e*., the shapes of the cutouts, and are robust to noise compared to other techniques (Figure 2a-e). LTSA, hLLE, mLLE and LDLE, which follow a bottom-up approach similar to RATS, failed to properly unravel the swiss roll datasets, exhibiting issues such as unnecessary folding, loss of aspect ratio, and significant boundary artifacts. These issues are also evident in the embeddings generated by the computationally intensive MVU, as well as its more efficient variant, lMVU.

**Figure 2.**
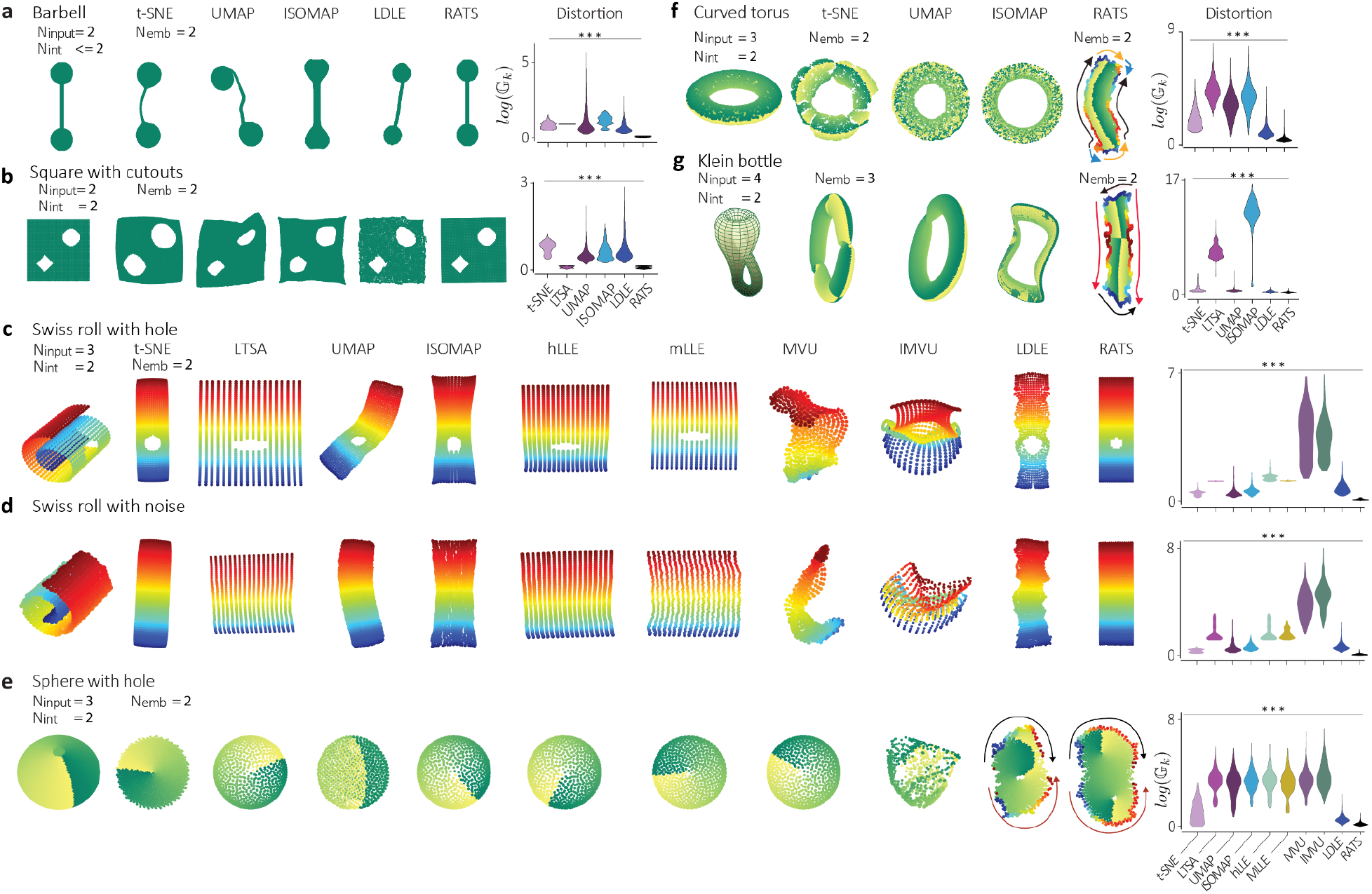
Distortion in embeddings of idealized datasets. a) Left: Barbell with N_int_ ≤ 2 embedded in N_emb_ = 2 using t-SNE, UMAP, ISOMAP, LDLE, and RATS, where N_input_ = 2. Right: log G_k_, Global distortion distributions for t-SNE, LTSA, UMAP, ISOMAP, LDLE, and RATS embeddings. b) Square with cutouts with N_input_ = 2 and N_int_ = 2, embedded using the same techniques as a). c) Embeddings (left) and global distortion distribution (right) for Swiss roll with circular cutout with N_input_ = 3 (N_int_ = 2) embedded in N_emb_ = 2 embedded using the same techniques as in a), and hLLE, mLLE, MVU and lMVU). d) Swiss roll with noise with N_input_ = 3 embedded in N_emb_ = 2, same as c). e): Sphere with a hole, N_input_ = 3 embedded in N_emb_ = 2, same as c). f) Left: Curved torus with N_input_ = 3 (N_int_ = 2) embedded in N_emb_ = 2 using t-SNE, UMAP, ISOMAP, and RATS. Right: Global distortion distributions for t-SNE, LTSA, UMAP, ISOMAP, LDLE, and RATS embeddings (Stats N = 5000 Kruskal-Wallis test in Supplementary table 1). g) Klein bottle with N_input_ = 4 (N_int_ = 2), here shown in 3D, embedded using the same techniques as f) with N_emb_ = 3 for all methods but LDLE and RATS which use N_emb_ = 2. (all statistics for these results using the Kruskal-Wallis test in Supplementary table 1).

**Table 1.** Hyperparameter range of the different methods.

| Method | Hyperparameters (all pairs are tested) |
| --- | --- |
| UMAP | n_neighbors in {10, 25, 50, 100, 200, 300}<br>min_dist in {0.1, 0.25, 0.5, 0.75, 1} |
| t-SNE | perplexity in {25, 50, 100, 200, 300, 500}<br>early_exaggeration in {8, 16, 32, 64, 128, 256} |
| ISOMAP | n_neighbors in {5, 10, 15, 20} |
| LTSA | n_neighbors in {5, 10, 25, 50, 100, 200, 300} |
| mLLE | n_neighbors in {5, 10, 25, 50, 100, 200, 300} |
| hLLE | n_neighbors in {5, 10, 25, 50, 100, 200, 300} |
| MVU | n_neighbors in {5, 10, 15, 20, 25} |
| IMVU | n_neighbors in {5, 10, 20, 30}<br>n_landmarks in {20, 40} |
| LDLE | k in {28, 21, 14}, eta_min in {5, 10}, gl_type in {"unnorm", "diffusion"} |
| RATS | k in {28, 21, 14}, eta_min in {5, 10}, cost_fn in {"distortion", "alignment_error"} |

Closed manifolds, such as spheres or tori, have no boundaries, and despite their intrinsic dimension being two, they are often processed in three or higher dimensions. RATS provides us with the ability to tear these manifolds and embed them into their intrinsic dimension. To study this, we first examine an idealized sphere with a hole (Figure 2e) and a toroidal dataset (Figure 2f), and compare the optimized embeddings from various methods. We find that when we embed in intrinsic dimensions (*N*_int_ = 2) using conventional methods, many of the methods yield collapsed embeddings for both datasets, consequently incurring high distortion (Figure 2e,f, Supplementary table 1). In contrast, RATS is able to provide a torn embedding whose gluing diagram indicates a toroidal and spherical topology, respectively (Figure 2E,F). While tearing closed manifolds can exacerbate distortion, as suggested by classical results in differential geometry such as *Theorema Egregium*^*47*^, RATS leverages tear information to compute tear-aware distances directly in the embedding space—without referencing distances in the original data space—yielding a low-distortion embedding (Supplementary table 1). Since other methods are unable to tear manifolds and provide collapsed embeddings, the question arises whether it is fair to compare these methods that cannot embed closed manifolds into intrinsic dimensions to RATS using distortion measures. To provide a more balanced comparison, we also evaluate embeddings of a curved torus and a ‘sphere with a hole’ by these methods in an embedding dimension (*N*_emb_ = 3) and compare them to the RATS embedding in their intrinsic dimension (Supplementary Figure 4a-b). Even under these circumstances, RATS can achieve comparable global distortion (Supplementary Figure 4b, full statistical outcomes in Supplementary table 1). Additionally, we show that the distortion calculated for RATS embeddings for increasing dimensionality “plateaus” at the true intrinsic dimensionality. This behavior results in a novel approach to estimate intrinsic dimensionality of the data manifold (see Methods, Supplementary figure 5a-b).

Computer vision applications, such as network topologies of images^1^ and conformation space of chemical molecules^48^, have uncovered non-orientable manifolds like Klein bottles. This raises the question of how manifold learning techniques respond to non-orientable data manifolds. To address this, we examined whether various techniques could embed a four-dimensional Klein bottle dataset into its intrinsic dimension, *N*_int_ = 2 (Figure 2g). Most manifold learning techniques are better equipped to embed these data into three dimensions, which results in a pinch or a self-intersection, leading to high distortion (Figure 2g). RATS, however, is able to embed the Klein bottle into its intrinsic dimension through tearing and provide a gluing diagram that correctly recovers its topology (Figure 2g). Among these methods, t-SNE, UMAP, ISOMAP, and RATS provided relatively low-distortion embeddings with RATS providing the one with minimum distortion (Statistics in Supplementary table 1).

### Evaluating RATS on biological datasets

Next, we examine whether biological data resembling closed manifolds can be described using the same manifold learning principles. We demonstrate that RATS can embed neural activity from both extracellular and calcium imaging recordings, and across both rodents and primates in a variety of brain regions and tasks where the underlying topology is often unknown or difficult to decode. We first turn to three recent studies in neuroscience that unveil intriguing topological manifolds in the neural population representations for head-direction^12,49^ and navigation systems^13,50^ in mammals.

In the first study, the authors examined neural activity from head direction cells in the anterodorsal thalamic nucleus (ADn) of a mouse while it foraged in an open 2D environment. They then embedded the 22-dimensional head direction activity dataset^12,49^, which exhibited a closed ring-like structure (Figure 3a). Although the intrinsic dimension of the ring is one, several conventional manifold learning techniques produced a representation of this topology^24^ that required at least two dimensions to produce a low-distortion parameterization of the data. To address this, the authors^12^ developed a specialized approach based on spline fitting to obtain a one-dimensional parameterization of the neural activity as it corresponds to heading direction. Here, we embedded the neural activity in *N*_emb_ = 2 dimensions using t-SNE, LTSA, UMAP, and ISOMAP and in *N*_emb_ =*N*_int_ = 1 dimension using RATS (Figure 3b), where the gluing instructions produced by the RATS embedding depicted a closed-loop topology, i.e. a ring. We demonstrate that decoding from a torn manifold in its intrinsic dimension offers advantages over decoding from a closed manifold in a higher-dimensional space. While we could directly decode the head direction angle from LDLE and RATS embeddings by simply scaling the one-dimensional embeddings to range between 0 and 2*π*, for other methods, we needed to employ the spline fitting approach^12^ on the two-dimensional embeddings to recover the intrinsic parameter (Figure 3c,d, Supplementary figure 6a-f). Thus, obtaining the intrinsic parameter directly is more generally applicable as it avoids the need to develop methods to estimate the parameter post-hoc from an intermediate-dimension embedding, i.e. in the case of closed topologies such as rings. We assess the mean squared error (MSE) between the decoded head direction angle from the embeddings and the angle measured by a light-emitting diode tracking (Figure 3c,d). The RATS embeddings result in the least error and these results generally hold with hyperparameter variation (Supplementary Figure 7).

**Figure 3:**
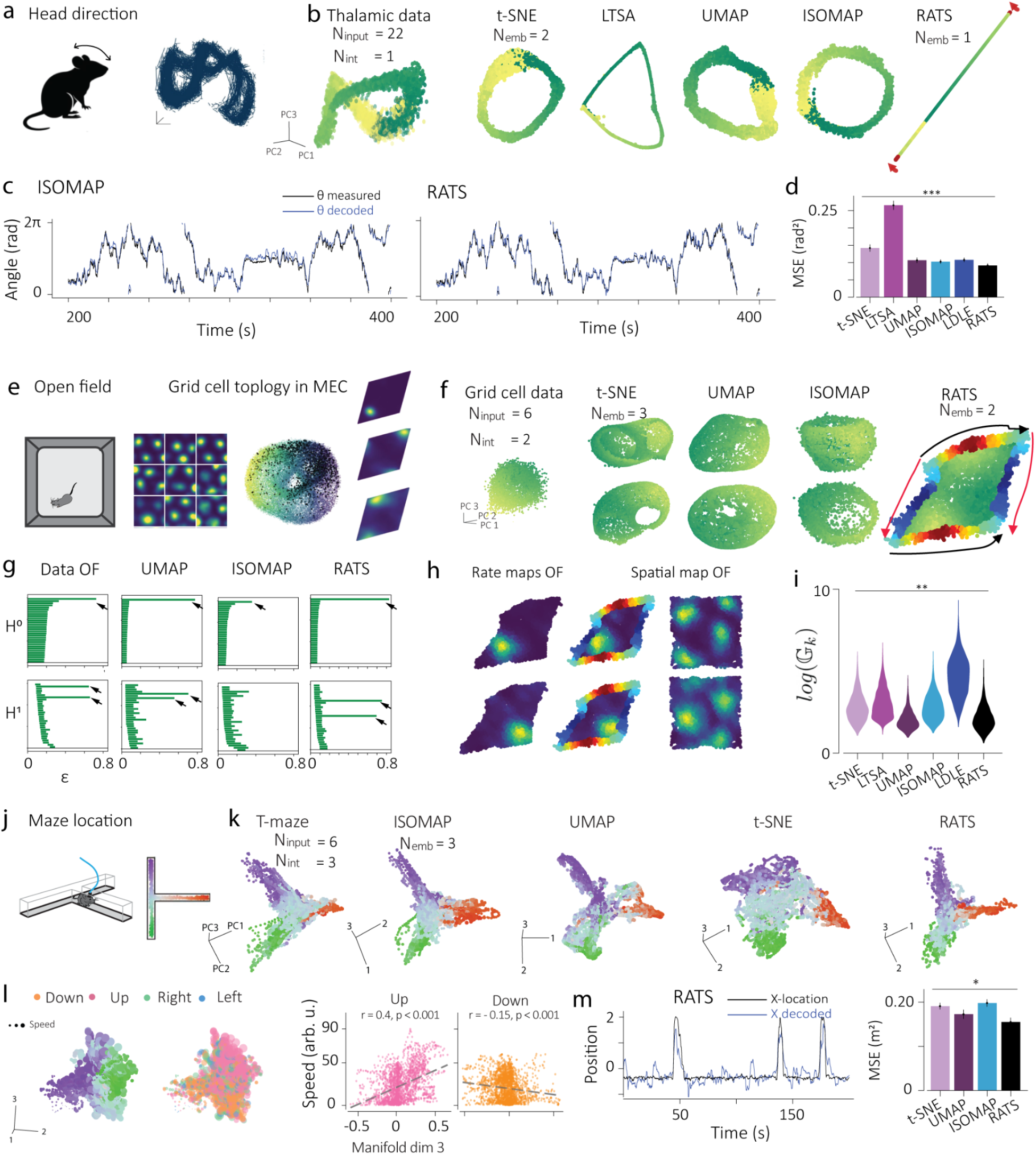
Neural activity from head direction cells, grid cells and place cells. a) The 22-dimensional population activity vectors recorded from ADn of mice while foraging in an open 2D environment have been shown to organize on a 1D manifold, visualized in 3D with ISOMAP^12^. b) PCA visualization of the input neural activity in 3D. Embeddings of the neural activity are obtained using t-SNE, LTSA, UMAP, and ISOMAP (N_emb_ = 2) and using RATS (N_emb_ =N_int_ = 1). RATS provides gluing instructions by coloring the ends of the 1D embedding, indicating a ring structure. c) Unsupervised decoding of the head direction angle from RATS and ISOMAP embeddings. d) Head orientation angle decoding MSE for all methods (Statistics in Supplementary Table 2). e) The 149-dimensional population activity vectors recorded from MECII of rats while foraging in an open 2D environment have been shown to organize on a toroidal manifold, visualized in 3D with UMAP^13^. f) Visualizing the input neural activity in 3D using PCA. Embeddings of the 6-dimensional PCA-reduced neural activity are obtained using t-SNE, LTSA, UMAP, and ISOMAP (N_emb_ = 3) and using RATS (N_emb_ =N_int_ = 2). RATS provides gluing instructions by coloring the two sides of the 2D embedding. The derived gluing diagram depicts toroidal topology (Figure 1E). g) The persistence barcodes for zeroth and first cohomology of the input data and the embeddings (see Methods), where ε denotes a scale parameter. h) Visualization of the firing rate maps of four of the grid cells on the RATS embedding and in the spatial coordinates. i) Global distortion distributions (Statistics in Supplementary Table 2). j) Illustration of T-maze adapted from ref^50^ that mice explored while Hippocampal CA1 CA^2+^ activity was recorded (N = 104). k) Left: Input data was obtained by dimensionality reduction using PCA (N_input_ = 6), visualized in N_emb_ = 3. Right: Embeddings in N_int_ = 3 using t-SNE, UMAP, ISOMAP, and RATS (also here in 3D). i) RATS embedding in 3D corresponding to the green and purple parts of the maze in j. Points are colored by the direction of movement and size corresponding to speed, showing that when in the green and purple zones of the maze, mice move faster upward (pink) and downward (orange), respectively. Left and right directions are also indicated in cyan and blue. Right: Correlation plots showing that the 3rd dimension of the manifold encodes information about movement speed in upward (pink, r = 0.4, p < 0.0001) and downward (orange, r = -0.15, p < 0.0001) directions. m) left: Linear decoding of position from RATS. Right: Position decoding MSE for all methods (Statistics in Supplementary table 2).

In other recent work^13^, authors investigated neural activity from grid cells in the medial entorhinal cortex (MECII) of rats, revealing that the activity patterns collectively reside on a toroidal manifold, with the rat’s movements in the environment corresponding to trajectories on the torus (Figure 3e). We compare manifold learning techniques on the 149-dimensional neuronal recordings reduced to six dimensions via principal component analysis (PCA; see methods for preprocessing of the data) to recapitulate the reported findings using UMAP with N_emb_ = 3 (Figure 3f). Since the intrinsic dimension of the toroidal object is two, we employ RATS to embed the data into its intrinsic dimensions. From the resulting gluing diagrams we identify a toroidal topology (Figure fF). We further validate the topology using persistence barcodes computed from the shortest path distances on the data and the embeddings (Figure 3g, Supplementary figure 8a). It is worth noting that only the tear information, *i*.*e*. the points along the tear, from the RATS embedding were utilized to compute the 1-homology, leading to a faster computation. The resulting persistence barcodes align with the ones derived directly from the data (Figure 3f). In fact, when visualized in 3D, the RATS tear depicts the two representative cocycles corresponding to a toroidal topology (Supplementary Figure 8b). We compared the spatial firing rate maps with the firing rate maps on the RATS embedding (Figure 3h, Supplementary figure 9a-f). We replicate these findings (Supplementary figure 10a-d) in an additional dataset from ^13^. We emphasize that RATS provides the means to directly embed in the intrinsic two dimensions with identified topology, whereas in previous work multiple steps were necessary to achieve the same results: extensive computation of cocycles was necessary to confirm the topology in the UMAP embedding and then a second embedding was carried out into intrinsic dimension for mapping the firing rates^13^. Depending on the application, the RATS embedding could thus complement Topological Data Analysis (TDA) algorithms that determine the homology of the data manifold to infer its topology. Comparing the lifetimes of the second and third persistent bars calculated for the different embedding methods reveals that RATS consistently recovers two persistent 1-cocycles across various hyperparameter settings (Supplementary Figure 11). Finally, we use local denoising to compute the global distortion metric for all embeddings and show that RATS provides the lowest distortion embedding (Figure 3i, Statistics in Supplementary table 2).

In another recent study^50^, calcium-imaging data of place cells from the rodent hippocampus underlying navigation of mazes with different geometries (T-shaped maze Figure 3j, Supplementary Figure 12a, and square maze, Supplementary Figure 13a) revealed structure that resembled the geometry of the mazes explored. Here we embed these data (N_input_ = 6) into the estimated intrinsic dimension of the data, which our analyses revealed to be N_int_ = 3 (Supplementary figure 12a). The third dimension represents the movement velocity of the mouse in the maze. We embed these data in N_int_ using t-SNE, UMAP, ISOMAP and also in 3D using RATS (Figure 3k) and find the optimal 2D projection of these embeddings for each method to linearly decode position. We find that RATS provides the lowest mean squared error for different location metrics (Figure 3l,m, Supplementary Figure 12b,c, Supplementary Figure 13b) for these data. Especially, for the T-maze, RATS is able to capture the structure of the maze more accurately than other methods (Figure 3k-m).

We, moreover, studied three neuroscience datasets where the neural population topology underlying tasks such as, sensorimotor control or decision-making, was complex, tangled, or unknown. In the first of these ^51^, we examine neural activity from the primary motor cortex of a rhesus macaque monkey^51^ during a manual cycling task where forward and backward pedaling was performed at different angular speeds to track a target (Figure 4a). Using linear dimensionality reduction methods, the authors reported that the neural population dynamics consisted of well-organized elliptical limit cycles in the first two principal components representing different speeds; yet the third dimension did not directly align with the different pedaling speeds (Figure 4b). We hypothesized that directly embedding the data into N_int_ = 2, using the tearing functionality of RATS, could recover a parametrization of both the cycling angle and speed. We embed neural activity, in *N*_emb_ = 3 dimensions using t-SNE, UMAP, and ISOMAP and in *N*_emb_ =*2* dimensions using RATS (Figure 4c), resulting in separate embeddings for each cycling direction. The gluing instructions produced by RATS depict a cylindrical topology, *i*.*e*. the manifold is closed along one axis and has boundary in the other. We find that these correspond to angle and speed, respectively.

**Figure 4.**
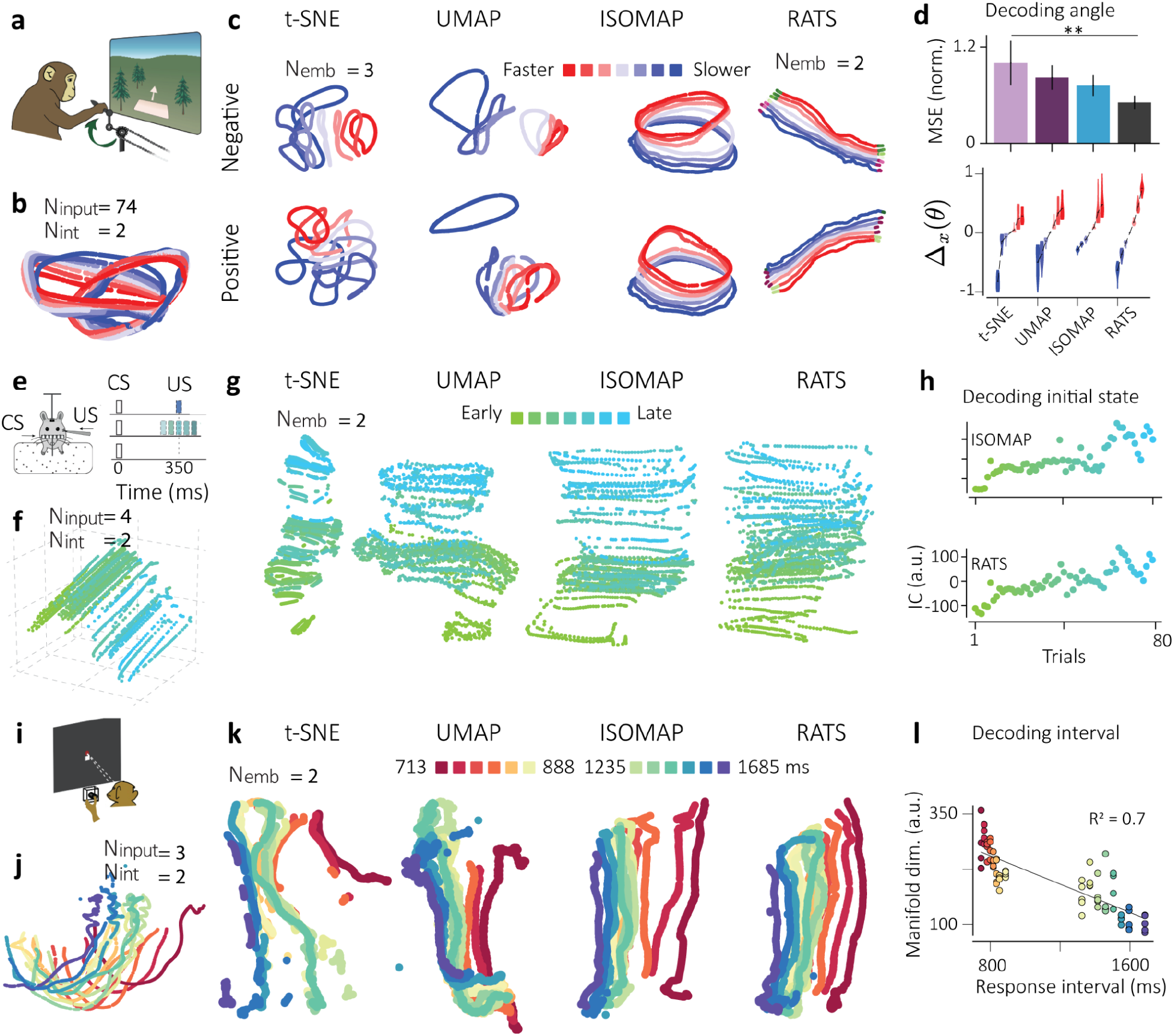
Embedding task-relevant neural activity from the motor cortex, cerebellum and thalamus in various cognitive and motor behaviors. a) Monkeys cycled forward (positive) or backwards (negative) at a range of speeds for juice reward while single units were recorded from the motor cortex. Adapted from ref^51^. b) Input data comprised of firing rates (N_input_ = 74) whose principal components are visualized here (N_emb_ = 3) suggest two interlocking ring topologies. Each line represents the conditioned-averaged response for each speed. Red to blue colors indicate different speeds. c) t-SNE, UMAP, ISOMAP and RATS embeddings for the 74-dimensional input data for the positive (bottom) and negative (top) pedaling directions. RATS embeddings represent a cylindrical topology in N_int_ = 2. d) top: Decoding of pedaling angle, bottom: distribution of angle-dependent distances between reference trajectory at speed bin 4 and each of the other speeds for positive direction (negative direction in Supplementary Figure 14c-f). e) Post-processed data from the Cerebellar Lobule IV/V, Simplex during a sensorimotor conditioning task ^52^ where mice were trained on temporal relationships between two sensory cues.f) Neural dynamics at the start of the trial exhibit state-dependent shifts, which when embedded using PCA (N_input_=4, N_emb_ = 3) reveals a nonlinear topology (N_int_ = 2). Green to blue colors show progression of time from early to late trials. g) t-SNE, UMAP, ISOMAP, and RATS embeddings for data in N_int_ = 2. h) Decoding of initial state along the manifold suggests a gradual linear drift ensues in the state during the course of the session. i) Context-dependent delayed motor response by monkeys during extracellular recording of thalamic single units ^54^. j) PCA (N_emb_ = 3) reveals a nonlinear manifold-like structure (N_int_ = 2). Red to blue colors indicate response times that span 640 - 1700 ms. k) Embeddings in N_int_ = 2 using t-SNE, UMAP, ISOMAP, and RATS L. Decoding along the manifold dimension recapitulates a strong correlation with the time interval generated by the monkey (R = 0.7, p <0.001). Statistics in Table 3

For each speed, we decode the pedal angle and assess the average mean squared error (Figure 4d, top), where decoding from the RATS embedding results in the least error. We next study how separated the trajectories corresponding to different speeds are in each embedding. If speed is independent of the angle in the embedding space then the distance between a pair of trajectories will be constant across all angles. Following the original work^51^, we chose an intermediate speed as a ‘reference’ trajectory, and computed the Euclidean distance to all other trajectories at every angle (Figure 4d bottom, alternative reference speed in Supplementary Figure 14a-f). RATS provides the smallest distance variance for all speeds with respect to the reference speed. Furthermore, for the RATS embedding, the speeds are close to being evenly spaced, as opposed to the other methods. Thus, RATS recovers a parametrization of the neural trajectories demonstrating that speed and pedaling angle are independent of each other, i.e. they disentangle in the motor cortex into separate orthogonal dimensions.

Following recent work^52^, we also examined neural dynamics of the rodent cerebellum (N = 75 single units, lobule IV/V) as mice learned the temporal relationship between a transient sensory stimulus (conditioned stimulus, CS) and a periocular airpuff (unconditioned stimulus, US)^52^. Having no a *priori* expectation about underlying topology in these data, we examined different task epochs before, during, and after behavior on each trial. We processed these data further using trial-by-trial firing-rate estimation using a variational autoencoder^53^ and reduce their dimensionality using principal component analysis to obtain input structure (Figure 4f, N_input_ = 4). Embedding the data obtained from before trial onset in apparent N_int_ = 2, we discovered a previously-unreported organization of initial condition (before the onset of the conditioned stimulus) appeared to form a continuous nonlinear topological manifold (Figure 4f). We hypothesized that the manifold dimension orthogonal to the dynamics during the time progression within a trial, represented drift in initial state before the trial started, which may relate to fatigue or state-dependent neuromodulation and may evolve in a linear manner. Of all the embeddings obtained from various methods, we find the RATS embedding to be most continuous and robust to distortion (Figure 4g). Several methods, including t-SNE, UMAP and RATS, provide decoding consistent with a linear assumption (Figure 4h, Statistics in Supplementary table 3).

Understanding latent variables that encode decision-making and cognition remains a challenge in neuroscience. We investigate data from the macaque thalamic nuclei during a decision-making task where monkeys used color and shape cues to determine whether they must respond with a certain effector and when^54^. Here, we uncover a novel nonlinear manifold in the neural activity (Nemb = 3 Figure 4J). When we embed these data into their intrinsic dimension (N_int_ = 2) using t-SNE, UMAP, ISOMAP and RATS (Figure 4k), we obtain a rectangular-shaped structure. Decoding along the dimension of the manifold that corresponds to the gradation, we obtain a systematic relationship between the manifold dimension and the average response time of the monkey. RATS provides the best quality of embedding for these data and the decoding of response interval is among the best, where ISOMAP also performs well in this case (Figure 4l, Supplementary Figure 15a-c, Statistics in Table 3). This provides evidence in favor of previous hypotheses^54^ that systematic increases or decreases in thalamic activity provide the cortex with contextual information by encoding corresponding variations in the animals’ production time.

### Evaluating RATS on manifolds generated by deep networks

We used deep recurrent networks to generate rich and organized topologies that resemble manifolds derived from the dynamics of previously-reported activity of neural populations^54–56^ and where we understand how latent variables along these manifolds relate to the networks’ behavioral outputs. Furthermore, we study scenarios where the manifold geodesics exhibit properties that can be expected from realistic datasets, such as, variable density, missing or noisy data, discontinuities and tearing of the manifold structure.

We first examine the influence of varying the density across the data points on manifold learning methods. We study the recurrent neural network dynamics underlying the task presented in Figure 4I, where the manifold was organized to reflect the logic of response times and the speed of the trajectory along the manifold was inversely related to the time interval generated by the network, resulting in a variation of densities within the manifold, which leads to warped embeddings from different manifold learning methods (Supplementary figure 16a-b). We found that for such data, RATS and ISOMAP provide the lowest distortion embeddings (Supplementary figure 16b-c). Noise is a frequently encountered nuisance variable in biological datasets^12,13,49,54^. To examine the robustness of manifold learning techniques to off-manifold and on-manifold perturbations, Gaussian noise was added to the input or to each network unit, resulting in exacerbated noise and density variations within and in the vicinity of the original data manifold.

In a second example, we recapitulate the findings of the cerebellar dataset using recurrent neural networks, this time to study how techniques perform under realistic variations in data such as, missing data, noise, frayed data segments, and high aspect ratios (Figure 4e-h). The network population activity during the associative learning task resembled an irregular Swiss roll-like object that varied in density, contained missing data segments, and was frayed at locations (Supplementary figure 16d-f). We found that while most techniques were able to embed these data manifolds into two dimensions (Supplementary figure 1e), they introduced significant distortions through tearing (t-SNE), non-uniform expansion (LTSA, UMAP), and overall warping (ISOMAP, LDLE). RATS is, however, able to preserve the discontinuities in the data manifold and provide the lowest distortion embedding among these (Supplementary figure 16f, Statistics in Supplementary table 4).

In addition to decoding metrics and providing data visualization, accurate low-distortion embeddings could enable the validation or invalidation of hypotheses in computational biology and other fields. Here we provide an example from neuroscience where recent proposals^17,57^ suggest that manifold-like topologies in neural population structure may enable different computations based on how information is decoded from the manifold, *i*.*e*., linear projections of the manifold embedded in high-dimensional space may serve some computations whereas, other computations may be more suited to readouts like geodesic distances along the manifold’s intrinsic dimensions. For example, recent work^55^ on time interval reproduction demonstrates that neural activity curves in semicircular trajectories (Figure 5a,b), potentially represent prior knowledge of time distributions that bias behavioral estimates in a manner consistent with Bayesian theory (Figure 5c). If prior knowledge is hypothesized to be encoded as curvature in such geometries, then unfolding such structures and decoding them along their intrinsic dimension should yield temporal estimates that are free of the influence of prior knowledge^57^, *i*.*e*., consistent with non-Bayesian models like Maximum likelihood estimates (Figure 5c). We find that manifolds representing such prior-related curvature, whose linear readouts are hypothesized to yield Bayes-like estimates (Figure 5c left), when embedded in lower dimensions and decoded, do indeed yield values that are consistent with Maximum likelihood estimation (Figure 5c, Supplementary figure 17a-b). Of the various techniques we tested, RATS provides the lowest distortion embeddings (Figure 5d,e). The quality of manifold decoding can often depend on features that are selected along intrinsic dimensions with different methods offering different trade-offs. We show that RATS consistently provides one of the lowest overall root mean squared error (RMSE) for different measures of decoding of time intervals from latent intrinsic variables (Figure 5d, Supplementary Figure 17c) and global distortion in its embedding (Figure 5e). Therefore, low-distortion embeddings can be used for hypothesis validation and for reliable decoding of relationships between latent low-dimensional variables and observable external variables.

**Figure 5.**
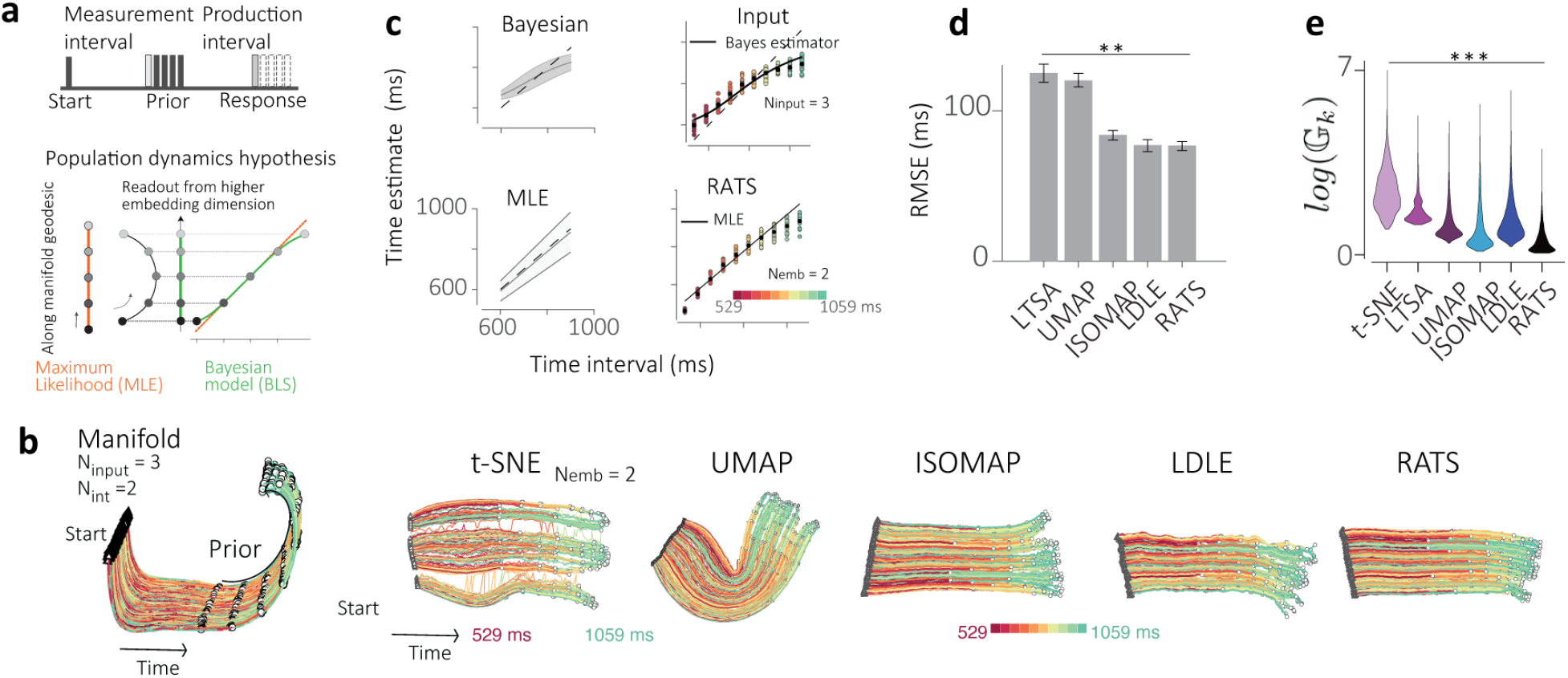
Hypothesis validation using manifold learning techniques. a) For a Bayesian interval reproduction task, temporal stimuli are sampled from a uniform prior distribution, which causes neural and network population trajectories to curve in a semicircular trajectory. Linear readouts of these populations lead to states that become compressed towards the mean of the prior, leading to biased estimates towards the mean of the prior, a hallmark of Bayesian inference using prior knowledge (BLS, green). Removal of the influence of the prior (orange), should lead to a mitigation of the bias but an increase in variability and the maximum likelihood estimator (MLE). b) Left: Nonlinear manifold formed by network dynamics in the temporal reproduction task exhibits time intervals along a semicircular curvature of the population manifold (N_input_ = 3). Right: Embeddings into N_int_ = 2 using t-SNE, UMAP, ISOMAP, LDLE and RATS. c) Left: Normative model of Bayes Least Squares estimation of time intervals and Maximum likelihood estimation. Right: Linear decoding yields estimates consistent with BLS. Linear decoding of the embedded manifold by RATS yields estimates consistent with MLE. d) Comparison of RMSE of decoded estimates from the stimuli ensuing from embeddings with different techniques. Error bars represent standard deviation. e) Global distortion distributions for the optimized embeddings provided by each technique.

## Discussion

With the emergence of low-dimensional topological and geometric structures in complex multivariate datasets across biological and non-biological fields, accurately isolating these structures into their intrinsic dimensions with minimal distortion to draw accurate scientific conclusions is crucial. To address this need, we introduced RATS, an unsupervised manifold learning technique capable of embedding complex datasets with topology, including closed and non-orientable manifolds, into intrinsic dimensions with consistently low distortion. Its robust performance across a variety of datasets, including real-world noisy biological datasets with unknown ground truth, establishes it as a versatile approach that especially mitigates the ubiquitous distortion problem commonly encountered in high-dimensional data science.

We have shown that RATS effectively embeds data in its intrinsic dimension, but an important question is how to determine that intrinsic dimensionality. Recent work^58^ has shown that linear approaches (e.g., PCA) overestimate the intrinsic dimensionality of neural data, whereas nonlinear methods provide a more accurate assessment. Similar to the use of variance explained to determine linear dimensionality, we demonstrated that our global distortion can be used in an effective, data-driven strategy for estimating intrinsic dimensionality: by embedding data in progressively higher dimensions until global distortion stabilizes.

The evaluation of embedding techniques is an open and debated question, amongst others most recently in bioinformatics for 2D visualization of, *e*.*g*., single-cell RNA sequencing data. Multiple measures have been proposed to quantify local or global structure^19,59–62^ and they often require supervised metrics like clustering accuracy or downstream task performance, which limits their generalizability. We introduced a novel measure of distortion, called global distortion, which quantifies geometric fidelity by comparing geodesics in the original space with those in the embedding space. In general, this is important for quantifying distortion when data are not embedded into Euclidean space, but into alternative geometries, *e*.*g*., by constraining the representation to lie on a sphere^31^ or in hyperbolic space, which is used for embedding hierarchical data. While estimating geodesics can be susceptible to “shortcuts” between points in noisy data, appropriate preprocessing mitigates this limitation (see methods, Figure 3I).

While none of the manifold learning methods presented here are optimized on our global distortion measure, our analysis confirms that different methods can indeed generate low-distortion embeddings. Furthermore, we demonstrate the practical utility of low-distortion embeddings in linking external variables to latent structures on the manifold and for validating hypotheses on biological computations. While we recognize that techniques like t-SNE and UMAP have shown to be powerful tools to visualize datasets, especially those where the clustering of data is highly informative, our findings highlight the implicit link between distortion in embeddings and analytical reliability, indicating that distortion reduction is critical for accurately interpreting how real-world variables are represented in latent manifold structures. As an important example of this, our experiments primarily focused on data whose parametrization of the underlying manifold was a physical quantity, *i*.*e*. space in the navigation circuit^13,12^ or time in motor control^54^. We both demonstrated that embeddings with high distortion had the highest decoding errors, as well as showed that RATS achieved the lowest distortion while incurring the lowest error in most cases.

By minimizing distortion, one can better reveal how real physical quantities are represented in latent low-dimensional representations of large datasets, since discrepancies in the representations of these quantities will correspond to real phenomena rather than artifacts introduced by the dimensionality reduction approach. Thus, quantifying distortion and reducing distortion in embeddings may be essential to distinguish whether given information is indeed absent from the data or whether an erroneous representation owing to distortion has masked our ability to access said information. It remains an interesting question how distortions affect latent representations of more abstract stimuli or complex behaviors of natural and artificial systems.

## Acknowledgements

The authors thank Juan Alvaro Gallego, Mehrdad Jazayeri, and Mikio Aoi for their comments on the manuscript. We thank Julius Koppen and Sowmya Narasimha for their help in processing datasets.

## Funding

This work was supported by funding from the Kavli Institute for Brain and Mind (DK), NIH (EB026936 to GM and DK), the NSF (CCF-2217058 to GM, EFRI 2223822 to GM and DK, DMS-2012266 to AC), a gift (Intel to AC), the Netherlands Organization for Scientific Research (Vidi-VI.193.076, Aspasia-015.016.012 and Gravitation-024.005.022 to DN), AiNed foundation (DN) and the NGF (NGF.1609.241.021 to DN).

## Author Contributions

DK, JSN, GM and DN conceived the study. All authors developed the analytical approach. DK and JSN implemented the methods. DK, JSN and KZ carried out all analyses. All authors wrote the manuscript. GM and DN supervised this project and provided funding for it.

## Competing Interests

The authors declare no conflicts of interest exist.

## Data and materials availability

Our generated datasets will be available at the conclusion of peer-review.

Our code will be made available at an open access repository at the conclusion of peer-review.

## Supplementary Materials

### Methods

#### Global distortion calculation

Let and *l*_*kj*_ be the shortest path distances from *x*_*k*_ to *x*_*j*_ in the data, and from *y*_*k*_ to *y*_*j*_ in the embedding, respectively. These are computed using the Dijkstra algorithm on the k-nearest neighbor graph on the ground truth (or denoised data) and on the embedding, respectively. Note that for the torn embedding we use the tear-aware shortest path distances as defined below. Then the global distortion at the *k* th point is given by the product of the worst-case expansion and the worst-case contraction

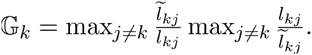

Finally, to measure the quality of a global embedding, we define its distortion to be a histogram built from global distortions at every point *i*.*e*. 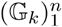.

#### Construction of k-nearest neighbor graph on the data and the embedding

To calculate shortest path distance for the global distortion calculation, we calculate k-nearest neighbor graphs on both the data and the embedding. The number of nearest neighbors, *k*, employed to construct the k-nearest neighbor graph is determined as the minimum value within the range of [9, 25] at which the distances of shortest paths on the data exhibit convergence. To ascertain this convergence, the following procedure is undertaken: The mean absolute relative change in the shortest path distances is computed for consecutive values of *k*. These are subjected to a threshold of 1e-3. The smallest value is identified for which the relative change remains below the tolerance for three consecutive values. In the rare situation where none of the value satisfies this condition, we relax the tolerance by doubling it, and repeat the process.

#### RATS Algorithm

Given *n* data points, the algorithm consists of three steps: (i) computation of *n* low-distortion local parameterizations that map the local neighborhoods on the dataset into lower dimensions, yielding a local view of the data. (ii) merge local views into *m* ≪ *n* intermediate parameterizations with larger neighborhoods, yielding intermediate views of the dataset. (iii) These views are aligned to obtain a global embedding by minimizing a quadratic over the product of orthogonal groups using Riemannian gradient descent. These steps are described in detail below.

### Local Parameterizations

Given a dataset 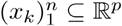 and an embedding dimension *d* ≤*p*, we start by computing a local neighborhood 𝒩_*k*_ around each point (using a nearest neighbor algorithm^63^). A local parameterization Φ_*k*_: ℝ^*p*^ → ℝ^*d*^, for the neighborhood 𝒩_*k*_ of *x*_*k*_ is then obtained by applying kernel PCA on the neighborhood *i*.*e*., on {*x*:*x* ∈ 𝒩_*k*_}, with an appropriate linear or nonlinear kernel *K*(·, ·). In particular, if *K* (x,y) = *ϕ* (*x*)^*T*^*ϕ* (*y*) for a certain choice of *ϕ*, then the *k*th local parameterization is given by Φ_*k*_ (*x*) = *P*_*k*_*ϕ*(*x*)*+ μ*_*k*_ where the *d* rows of *P*_*k*_ contain the *d* leading eigenvectors of the covariance of {*ϕ* (*x*) : *x* ∈ 𝒩_*k*_}, and *μ*_*k*_ represents the centroid of the same collection. In our experiments, we utilized the linear kernel *K*(*x,y*) = *x*^*T*^*y* so that *ϕ* (*x*) = *x* and the cosine kernel *K* (*x,y*) = *x*^*T*^*y/* ‖*x* ‖_2_ ‖ *y* ‖_2_ where *ϕ* (*x*) = *x* / ‖*x* ‖_2_. In practice, the explicit value of *ϕ*(*x*) is not needed since the calculations involve dot products that can be directly evaluated using the kernel function.

### Local distortion calculation

Local distortion is calculated as the product of maximum expansion and maximum contraction of pairwise distances from the high-dimensional data to the low-dimensional embedding within a local neighborhood, where we use Euclidean distances in both spaces. Let Ψ denote a generic local parameterization on a set of points 𝒰, local distortion is defined as,

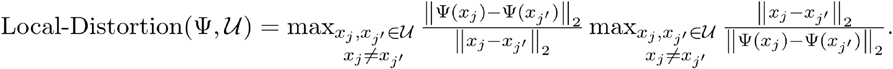

Then the distortion incurred by the *k*th local parameterization on the *k* ′ th neighborhood is obtained by substituting Φ_k_ for Ψ and 𝒩_*k*_*′* for 𝒰 *i*.*e*. Local-Distortion(Φ_k_, 𝒩_*k*_*′*.

### Postprocessing Local Parameterizations

To remove anomalous local parameterizations which incur abnormally high local distortion, a postprocessing step^27^ may be performed by repeatedly replacing a parameterization Φ_*k*_ of the neighborhood 𝒩_*k*_ of *x*_*k′*_ with that of a neighbor Φ_*k*_′ where *x*_*k′*_ ∈ 𝒩_*k*_, that produces the least local distortion on 𝒩_*k*_ *i*.*e*.,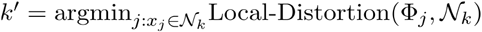. This procedure is repeated until no further replacement is possible.

### Intermediate Parameterizations and Views

Each of the *n* local parameterizations Φ_*k*_ yields a local view Φ_*k*_ (𝒩_*k*_ = {Φ_*k*_(*x*):*x*∈ 𝒩_*k*_}. A large number of small local views lead to high computational cost and instability during their global alignment^27^, due to small overlap between the small neighborhoods^64^. Therefore, through a clustering procedure we obtain *m* ≪ *n* intermediate neighborhoods 𝒱_*i*_ yielding *m* intermediate views 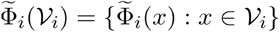. At any given stage of the procedure, the points in the *i*th cluster are denoted by 𝒞_*i*_ and the cluster label of the *k*th point is given by *c*_*k*_. The intermediate neighborhood 𝒱_*i*_ associated with the *i*th intermediate view 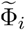 is defined to be the union of the neighbors of the points in cluster 𝒞_*i*_ *i*.*e*., 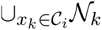. Beginning with each point belonging to its own cluster with an associated parameterization (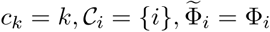 for *k,i* to *n*), the clustering procedure moves points into clusters based on the following rules: (i) a point can only move into one of its neighbors’ clusters, (ii) it cannot move into a cluster that is smaller than the cluster it currently belongs to, and (iii) among the remaining potential clusters it will move to the one whose parameterization results in the least cost, where the cost can be defined in multiple ways. The cost of moving the *k*th point into the *i*th cluster is based on the local distortion of the *i*th cluster’s parameterization 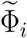 over the union of the cluster’s neighborhood 𝒱_*i*_ and the local neighborhood of the point 𝒩_*k*_ or based on the alignment error on the points in the overlap between the cluster’s neighborhood and the neighborhood of the point i.e. the alignment error between 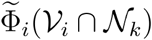 and 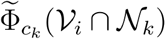. This procedure is a greedy iterative procedure to merge the points into clusters until the minimum number of points in each cluster reach a user specified threshold. After pruning the empty clusters, the parameterizations associated with the remaining clusters are denoted 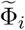.

### Aligning Intermediate Views for a Global Embedding using Riemannian Gradient Descent

To obtain a low-distortion global embedding, we find rigid alignments between the intermediate views. For convenience, we define the parametrization of the *k*th point in the *i*th intermediate view as 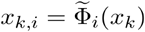 and, let Г be a bipartite graph between *n* points and *m* views. An edge Г in is given by a tuple i.e. (k,i) ∈ *E*(Г) if *x*_*k*_ ∈ 𝒱_*i*_. Then the aim of rigid alignment is to find *m* orthogonal matrices 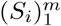 and translation vectors 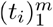, that minimize the alignment error, 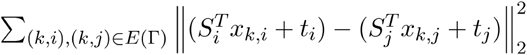 so that on average after alignment, the parametrizations of a point in two views containing the point are as close as possible.

An equivalent problem is

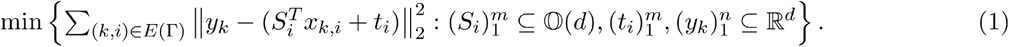

Following further reformulation of this problem as a quadratic program on the orthogonal matrices 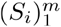 ^6564^. As opposed to the affine alignment of local views, which results in an optimization problem with a closed-form solution, the choice of aligning the local views using rigid transformations results in a quadratic program on orthogonal matrices that corresponds to a non-convex optimization problem without a closed-form solution. Despite this added complexity, our experiments demonstrate that rigid alignment yields embeddings with substantially lower distortion than those obtained via affine alignment. Moreover, while non-convex optimization methods like gradient descent are typically prone to convergence to suboptimal local minima, our approach is supported by recent theoretical results^64^ that guarantee local linear convergence and noise stability for Riemannian gradient descent applied to this specific objective.

Note that LTSA uses *affine* transformations for global alignment and assumes that the feature vectors are uniformly distributed in the embedding *i*.*e*., *YY*^*T*^ *= I*_*d*_ where *Y* = [*y*_1_, …,*y*_*n*_]. The uniformity on the embedding is also assumed by mLLE and hLLE. As a result, the aspect ratio of the data manifold is not preserved in the embeddings produced by these methods. In contrast, we use rigid transformations for alignment with no assumption on *Y*, preserving the aspect ratio and thus incurring less distortion.

### Tearing data manifolds

To embed a data manifold that is either closed, non-orientable or exhibits high curvature, into the intrinsic dimension, the manifold must be torn apart (as a cartographer flattens the three-dimensional earth into a two-dimensional map). We achieve this by first constructing an over-torn embedding, which is then progredssively stitched back together through an iterative procedure. The over-torn embedding is generated in two steps: we begin by extracting a spanning tree from a graph where nodes represent intermediate views and edges are weighted by the rank of the overlaps between them. Next, each child view is aligned to its parent using a single instance of the orthogonal Procrustes problem. The embedding is then refined and stitched together through the subsequent iterative process. At every iteration, we first compute local neighborhoods 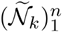 in the embedding. Then the *ii*th intermediate neighborhood 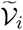 in the embedding is the union of the neighbors of the points in cluster 𝒞_*i*_, *i*.*e*. 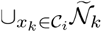. Note that these may differ from the intermediate neighborhoods in the data 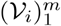. Given these intermediate neighborhoods in the data and the embedding, instead of aligning all overlapping intermediate views we align *only* those views that are overlapping in both the data (𝒱 _*i*_∩ 𝒱_*j*_≠ Ø) and the embedding (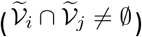) *i*.*e*., we solve for orthogonal matrices 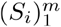 and translation vectors 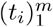, that minimize the alignment error,

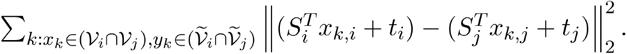

In particular, we define a bipartite graph Г ′ such that *V*(Г ′) = V(Г) while (*k,i*) ∈ *E* (Г ′) if *x*_*k*_ ∈ 𝒱_*i*_ and 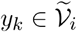. Finally, we solve Eq. (1) by replacing Γ with Γ′ throughout. In each iteration, the alignment results in increased overlap between intermediate neighborhoods in the embedding. At the end of the procedure, for manifolds such as a sphere or a Kléin bottle, a tear is retained, while for others, such as a swiss roll or barbell, the tear vanishes as the intermediate views are fully stitched together.

### Gluing instructions for the torn embedding

To visually represent points that are in close proximity in *y*_*l*_ ∼ *y*_*l*_ ^*′*^ the data space but appear on opposite sides of the tear, we color their embeddings with a similar color.

We say that the embedded points *y*_*l*_ ∼ *y*_*l*_ ^*′*^ if the following conditions are satisfied

1. The points *x*_*l*_ and *x*_*l*_ ^*′*^ lie in different clusters *c*_*l*_ ≠*c*_*l*_
2. The points *x*_*l*_ and *x*_*l*_ ^*′*^ are in the intersection of the corresponding intermediate views in the data i.e. 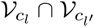
3. The embeddings *y*_*l*_ and *y*_*l*_ are far apart as captured by the empty intersection of the corresponding views in the embedding 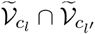.

A graph on the embedded points where edges correspond to the relation *y*_*l*_ ∼ *y*_*l*_ ^*′*^ is constructed. The coloring of the tear is obtained as the eigenvectors of the Laplacian of the above graph. By tracking the corresponding colors along the tear, we obtain “gluing” instructions for the torn embedding to recover the topology of the data manifold (Figure 1d, Supplementary figure 2a-b).

The resulting torn RATS embeddings for closed manifolds may have rough edges that can be mitigated by adjusting the tear-positioning using a semi-automated cut-and-paste method within a graphical user interface (Supplementary Figure 2a-b). This process maintains the same topology while producing an embedding with a smoother tear.

### Tear-aware Shortest Path Distance

Let *d*_0_(*y*_*k*_, *y*_*k*_) be the naive shortest path distance between *y*_*k*_ and *y*_*k*_ ^*′*^ computed using Dijkstra algorithm on the k-nearest neighbor graph on the embedding (k=10 in all of our experiments). If *y*_*l*_ ∼ *y*_*l*_ ^*′*^ then, by definition, the two points have local representations due to both the *c*_*l*_th view (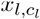 and 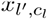) and the *c*_*l*_^*′*^th view (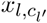 and 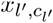). Then we define the -hop shortest path distance *d*_*t* − 1_(*yk,yk ′*) as the minimum *d*_*t*_(*yk,yk ′*)of and

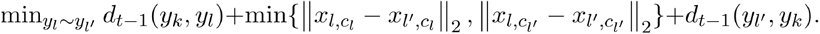

The tear-aware shortest path distance 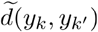 is obtained by computing *d*_*t*_ for increasing values of *t* ∈ {1, …,*t*_*max*_} until the convergence of *d*_*t*_. We use *t*_*max*_ = 5 in all of our experiments.

### Comparison to other bottom-up manifold learning techniques

RATS belongs to a family of bottom-up manifold learning techniques, which first construct local parameterization of the data, i.e. every point has its own mapping so that its local neighborhood has a low-dimensional and a low-distortion representation. The methods, including LTSA, hLLE and RATS use a form of local PCA for local views, where RATS also allows for local kernel PCA (which enables non-Euclidean similarity metrics) to construct locally nonlinear views. In contrast, LDLE uses a spectral construction procedure that selects (for each neighborhood) subsets of global eigenvectors of the graph Laplacian representation of the data. Due to the oscillatory nature of the Laplacian eigenvectors and their degeneracy near the boundary of the data manifold, the local views constructed in LDLE can incur high local distortion.

The local views are then aligned together to obtain a global embedding. The aim of the global alignment step in these methods is to find transformations of each local parametrization, so that the representations of the overlap between neighborhoods, due to the respective (transformed) parameterizations, are as close to one another as possible. This corresponds to minimizing the alignment error between transformed local views on the overlapping points. In LTSA, the global alignment step assumes a uniform distribution on feature vectors in the embedding space which allows the global transforms to be affine, thus limiting its capacity to preserve the aspect ratio of the manifold, and often leading to highly-distorted embeddings. The same uniformity assumption is also made by mLLE and hLLE, likewise preventing their ability to maintain the aspect ratio in their embeddings. On the other hand, RATS makes no such assumptions about the data, and uses rigid transforms instead for the alignment, which preserves distances between points, and is key to its achieving low-distortion embeddings. RATS utilizes Riemannian gradient descent to minimize the alignment error with local linear convergence and noise stability guarantees^64^.

LDLE similarly uses rigid transforms. However it uses a heuristic Procrustes-based method to calculate the global alignment, which has two drawbacks: it has slow convergence, and it fixes misalignments locally, i.e., converges to a suboptimal local minimum. This in addition to its local view construction, leads to higher distortion than RATS.

### Manifold denoising for distortion calculation

While the low dimensional embeddings are computed from the noisy data, they are intended to reflect the clean geometry. Therefore, directly evaluating the distortion of an embedding relative to the noisy data can be unreliable. Global distortion calculated given noisy data and an embedding derived from that data can be seen as an upper bound on the the global distortion calculated given the unobserved clean data and an embedding derived from the noisy data, i.e. the latter being a measure of whether the embedding derived from noisy data preserve the global geometry of the clean data.

To address this, we denoise the data as a more stable reference for assessing distortion. Specifically, we apply the local linear denoising method^38^ as a preprocessing step to calculating geodesic distances in the high-dimensional data for the distortion calculation. This approach leverages the assumption that, within a small neighborhood, the data can be well-approximated by a linear subspace. By projecting the data onto these local subspaces, the method filters out noise, and the denoised neighborhoods are then combined through mean averaging and a smoothing procedure^38^.

### Hyperparameter optimization for manifold learning techniques

(UMAP, ISOMAP, t-SNE, LTSA, mLLE, hLLE, MVU, IMVU, LDLE, RATS). We performed a grid search optimization based on the principal hyperparameters of each method. The optimal embedding was based on the hyperparameter set that yielded the minimal value for the maximum of the pointwise global distortions, i.e.. The following table shows the ranges of hyperparameters tested during the grid search optimization for each of the competing methods:

### Kruskal-Wallis Statistics

Given multiple groups (manifold learning algorithms) with non-normal distribution statistics, we administer a non-parametric Kruskal-Wallis test throughout to consider the null hypothesis that the pointwise global distortion due to each algorithm (groups) originates from the same distribution. The test is implemented using the scipy.stats.kruskal method from the scipy Python package with default hyperparameters to obtain the H statistic and the p-value. At a significance criterion of 1e-3, the null hypothesis is rejected across all the datasets considered in this work.

### Two-sided Wilcoxon Test Statistic

To test the null hypothesis that the pointwise global distortion due to the embeddings generated by two algorithms originates from the same distribution, we employ the two-sided Wilcoxon signed-rank test. The test is implemented using scipy.stats.wilcoxon method from the scipy Python package with default hyperparameters to obtain the W statistic and the p-value, and a significance level of 1e-3 is used throughout.

### Identifying intrinsic dimension

We embed the data using RATS into progressively higher dimensions. For every dimensionality we calculate and plot the global distortion of the resulting embedding. As with cumulative variance explained for PCA and Kullback–Leibler (KL) divergence for t-SNE, the “knee” of this plot provides an estimate for the nonlinear dimension of the manifold. In other words, the smallest dimension at which the (log-)distortion plateaus serves as an upper bound for the intrinsic dimension of the data manifold. We demonstrate this for flat manifolds where the aforementioned dimension accurately corresponds to the intrinsic dimension (see e.g., flat Klein bottle Supplementary figure 5b) and for the curved manifolds (e.g., a hypersphere, Supplementary figure 5a).

### Head direction cell population dataset

We took the spiking activity of 22 neurons in the anterodorsal thalamic nucleus (ADn) region of a mouse that was awake and foraging in an open environment along variable paths with variable velocities ^12^. The spike activity is preprocessed in exactly the same manner as in ^12^: firing rates are first computed by convolving the spike times with a Gaussian kernel of standard deviation 100 ms. Then the rates are replaced by their square root to stabilize the variance. Finally, the first 5000 rates form the dataset in our analysis. We embedded this 22-dimensional dataset using various manifold learning techniques.

### Grid cell population datasets

We took the simultaneously recorded extracellular spikes from the 149 “pure” grid cells identified by ^13^ from the MECII region of a rat’s brain while it forages in an open field. Similar to their preprocessing procedure for the topological analysis of this data, we preprocessed the neural activity by first applying Gaussian smoothing with a window size of 50ms, then selecting the 15000 most active population vectors which is then followed by a PCA projection into 6 dimensions. We follow the downsampling technique used in^13^, which is based on a point-cloud density strategy motivated by a topological denoising technique^66^ and a fuzzy topological representation used in UMAP^24^. For downsampling we use a cosine metric and a neighborhood size of 1500, and obtain a topologically denoised subsample of size 4800. Finally, we embedded this 6-dimensional dataset using various manifold learning techniques. To be consistent with the analysis in the original study^13^, we use the cosine metric with all techniques and specifically, we use kernel PCA with the cosine kernel to construct local views with RATS. Additionally, while LTSA is limited to linear-local views of the data, RATS adopts a more flexible approach, allowing the utilization of kernel techniques for modeling the data locally.

The firing rate maps of the grid cells are visualized in spatial coordinates and on the smoothed RATS embedding (Supplementary Figures 9 and 10D).

Similar to our distortion calculations, for the persistent homology analysis, we also constructed *k*-nearest neighbor graphs. Using these, we computed the shortest-path distances between all pairs of points in the data and their embeddings. For the RATS and LDLE embeddings, we utilized tear-aware shortest path distances between the subset of points that lie on the tear. Finally, we calculated the persistence barcodes for the zeroth and first cohomology by applying RIPSER^67^ to the shortest path distance matrices (Supplementary Figure 8a).

### Maze exploration dataset

We analyzed calcium imaging data from the CA1 region of the mouse hippocampus during exploration of a maze^50^. The calcium imaging data for the T-Maze(N = 104 units) was preprocessed by applying PCA to the activity matrix, reducing it to N_input_ = 6. These data were embedded into N_int_ = 3 using t-SNE, UMAP, ISOMAP, and RATS (Figure 3k). For RATS we used kernel PCA with a radial basis function (RBF) kernel. For each embedding, we optimized projection angles into two dimensions to align to the physical maze coordinates using linear regression, and calculated MSE. We colored points in the embedding based on their combined mean squared error relative to a global threshold (95th percentile across all methods). Points exceeding this threshold were highlighted to identify regions of poor alignment within each embedding.

To further explore the structure of the three-dimensional RATS embeddings, we colored the embedding according to the animal’s movement direction (upward or downward) at each timepoint, with point size reflecting instantaneous movement speed (Figure 3l). To quantify the distribution of movement directions along this dimension, speed of the animals were correlated with their manifold dimension 3 values (Figure 3L) and Pearson’s correlation coefficient was determined. To visualize how embedding values along manifold dimension 3 related to the four movement directions, we plotted the points using polar scatter plots (Supplementary Figure 11) where points were grouped by movement velocity: direction ‘up’ and ‘down’ points were plotted along a vertical axis, and ‘left’ and ‘right’ points along a horizontal axis. Points were slightly jittered and plotted in random order. Points were colored according to the animal’s movement direction at each timepoint, with point size reflecting instantaneous movement speed. The radial position of each point reflected its embedding value on the selected dimension (after adjusting for polarity and centering based on the median of opposing directions). Separate polar plots were generated for each embedding method (Isomap, UMAP, RATS) to compare the distribution of embedding values relative to movement velocity. For RATS and Isomap the third dimension encodes movement in the vertical direction of the maze, while there is no apparent organization of movement along the horizontal direction.

We additionally analyzed calcium imaging data obtained during exploration of a square maze. The calcium imaging data for the square maze (N = 115 units) was preprocessed by applying PCA to the activity matrix, reducing it to N_input_ = 10. This 10-dimensional dataset was embedded into N_int_ = 2 dimensions using t-SNE, UMAP, ISOMAP, and RATS. We aligned the embeddings to the two dimensional physical coordinates, by solving an instance of the orthogonal Procrustes problem, and calculated MSE in X and Y coordinates (Supplementary Figure 13b).

### Motor cycling dataset

We analyzed preprocessed recordings from 74 neurons in the primary motor cortex of a monkey performing a cycling task in a virtual environment^51^. The monkey controlled movement by hand-pedaling a cylindrical device either forward or backward at fixed target speeds labeled 2 through 8 (∼0.8–2.1 Hz). After uniformly downsampling the data by half, we obtained 7051 population activity vectors. Constructing a nearest neighbor graph (e.g., with 35–45 neighbors) on this dataset yielded 14 disconnected components, each corresponding to a unique combination of cycling direction and speed. To reveal organization along the speed axis in the embedding space, we added edges from each population activity vector to its nearest neighbors from all other cycling speeds. This resulted in a nearest neighbor graph with two connected components—one for the positive and one for the negative direction. This indicates that the neural activity is more similar across speeds within a cycling direction than for the same speed across cycling directions. We then computed two-dimensional *torn* embeddings using RATS and three-dimensional embeddings using other manifold learning techniques.

To decode the pedal’s angular coordinate from the RATS embedding, we processed the 2D embedding for each cycling speed and direction by rotating the embedding so that the first principal component aligns with the x-axis, and then scaled the resulting x-values to lie between and. For the other methods, we extracted one-dimensional angular coordinates from 3D embeddings, corresponding to a given cycling speed and direction, using a spline-fitting approach^12^. We computed the mean squared error (MSE) between the decoded and true angular coordinates for each method. Among all methods, the RATS embedding consistently achieved the lowest MSE (Figure 4D).

Following^51^, we investigated how target speed relates to the structure of embedded trajectories. We start by fixing a cycling direction and selecting a reference trajectory *T*_ref_(*θ*), along with a set of equally spaced phase angles *θ*_i_ = 10 + 20*i* (in degrees) for *i* = 0 to 17. For each *θ*_*i*_, we compute the distance between the reference and all other trajectories having the same cycling direction. Specifically, for a test trajectory *T*_ref_(*θ*), the distance at *θ*_*i*_ is defined as 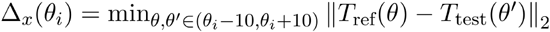. We visualize these trajectory differences using violin plots (Figure 4D, Supplementary Figure 14c-f). For the RATS embeddings, the relationship between the median distances and target speed exhibits a linear trend, while the other methods displayed nonlinear patterns.

### Trace conditioning dataset

Mice performed a trace conditioning task while extracellular recordings were performed in Lobule IV-V/Simplex of the cerebellar cortex. Mice learned an association between a conditioned stimulus and an unconditioned stimulus. Single units were isolated (N = 185) and processed using a variational autoencoder to obtain firing rate estimates^53^ for each trial within sessions for a single individual. Multiplexing the firing rates from three separate sessions, based on their dynamical similarity^53,68^, and after controlling for density variations, PCA was performed to denoise and reduce the dimensionality of the data from 75 to 3 dimensions (using a cumulative explained variance threshold of 65%). Neural trajectories from 300 ms before the onset of the conditioned stimulus were studied (N = 80 samples). This initial period before the onset of the trial revealed a continuous organization orthogonal to the dimension in which the time-varying dynamics association with the task evolved, giving rise to a nonlinear manifold (N_emb_ = 3, N_int_ = 2). These data were embedded into N_int_ = 2 using various manifold learning methods and subsequently aligned using rigid transformations (Figure 4E-H). The axis orthogonal to the time-varying dynamics during trials was decoded (Manifold dim 2) and position along this dimension was plotted against trial number to reveal a near-linear progression in initial state. We model the drift in the initial state as a linear process and calculate mean squared error with respect to a cross-validated linear model on the data.

### Motor timing dataset

In the motor timing task from ref^54^, monkeys were trained to respond to color cues (Cue) to generate time intervals of different durations. Figure 4I illustrates the task structure, where after the presentation of the Cue, a transient Start flash is presented (Start). The monkey was trained to generate different time intervals by a button press or a saccadic eye movement (Response). This led to the generation of response intervals (Tp) from the onset of the Start flash that depended on the cued context. The error feedback with respect to the desired time intervals (Ts) was provided by enforcing a time window within which responses were rewarded. The study^54^ also put forth a hypothesis that the contextual signal, encoding context-dependent timing, was supplied to the cerebral cortex via step-like inputs of graded magnitude from thalamic neurons. However, this hypothesis remained untested.

We analyzed the single unit extracellular recordings from the thalamus of a single animal (N = 402 units), across 20 sessions, when the subject used the hand as effector. We divided the successful trials into eight bins for the short and long interval condition each based on the response time and calculated the PSTH for each bin from 75 ms after the Start flash to the maximum of the longest timebin. We downsample this data N = 6774 points and apply density correction before applying PCAto reduce the dimensionality of the dataset to Nemb = 3(using a threshold of 45% based on cumulative explained variance). Visualization revealed a nonlinear manifold with gradation according to response interval that was consistent with the predicted hypothesis (Figure 4j). To decode response interval and test the hypothesis, we embedded these data in apparent N_int_ = 2 using t-SNE, UMAP, ISOMAP and RATS (Figure 4k). Decoding was obtained by performing rigid alignment corrections to all embeddings and obtaining the manifold dimension orthogonal to the time course of the trial and examined the correlation with response time for each embedding (Figure lL, Supplementary Figure 15a-c, Statistics in Table 3).

### Network training, tasks, and manifold generation

Idealized datasets provide validation of ground-truth manifolds, whereas in real-world datasets, often the ground-truth is unknown. Under these circumstances, neural network models can provide a means to generate large-scale datasets whose underpinnings obey manifold-like topologies, which are also riddled with realistic features, such as, uneven density, noise, discontinuities. In this work we also develop methods to control and manipulate noise along and off the manifold, as well as vary the density of data points across the manifold. This allows us to investigate the impact of these features on distortion using different techniques.

Our examples are drawn from the field of neuroscience, where Recurrent Neural network models (RNNs) have demonstrated remarkable success in recapitulating neural population activity^69–72^. Specifically, we utilize topologies generated by the dynamics of neural networks representing three neuroscientific behavioral tasks ^54–56^. Each of these networks emulates the properties and encoding principles of neural population dynamics believed to underlie various behaviors in the macaque cerebral cortex and the mouse cerebellum. By employing RNNs, we not only recreate the dynamics underlying these tasks but discover families of population trajectories that remain confined to manifold-like geometries with quantifiable intrinsic dimensions.

### Motor Timing network datasets

Following the findings of the previous work^54^, as described in the section on the Motor Timing dataset, the RNN implementation assumes that the recurrent connectivity models the cortical region responsible for encoding time production signals. Additionally, a time-dependent step input is used to model the subcortical contextual cue, whose magnitude sets the time interval production dynamics in the cortex.

The RNN consisted of *N*=200 nonlinear units with a hyperbolic tangent activation function. Let the activity of the population be represented by the time-varying vector x (*t*), whereas the firing rate of such a population be vector r (*t*). We assume that the time constant of each unit τ is 10 ms. Each unit also contains a bias term (*c*_*x*_) and a term for internal noise (drawn from 𝒩(0,0.01), which was sampled at each time step (*δt* = 1 ms). The Start pulse is administered at a random time after the presentation of the contextual Cue input to the network. The target function was a ramp specified after the Start until the network output crossed a threshold at the desired time interval for each trial *t*_*s*_. The actual time at which the network crossed the threshold (Response) was taken as production time for a given trial *t*_*p*_.

The network dynamics were governed as per the following equations:

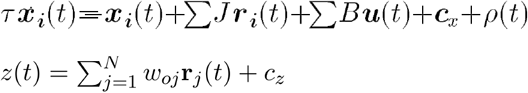

The recurrent weight connectivity matrix *J* was initialized by drawing values from a normal distribution with zero mean and variance 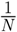^73^. The network received a two-dimensional input **u**(*t*) consisting of a context cue *u*_*c*_(*t*) and a transient pulse at a later time, *u*_*s*_(*t*), which marked the time from which the network generated the desired time interval. Model parameters, *J,B,w*_*o*_,*C*, were trained using backpropagation through time^74^ by minimizing a squared loss function between the network output *z*(*t*) and a target function, which was a linear ramp starting at the time of Start and crossing a threshold at the desired time for a given context input.

### Trace conditioning network datasets

In the second task of *Trace conditioning*, mice learned to associate between two sensory cues, viz., a brief flash of light (Light) and a brief periocular stimulation (Airpuff) after a fixed delayed interval. This behavior is thought to be encoded in cerebellar cortical circuitry via basis set inputs from cerebellar granule cells. The second source of the input signaling sensory events arrives at the cerebellar cortex via the climbing fibers. To mimic this input regime carrying sensory and temporal information, RNNs received multi-dimensional time-varying input *u*(*t*) through synaptic weights B =[*b*_0_,*b*_*g*_*c,b*_*c*_*f*]. The input *u*(*t*) included a step input, *b*_0_, the summation of decaying Gaussian granule cell basis kernels *b*_*g*_*c* for the Light and Airpuff, and a 10 ms pulse to mark climbing fiber activation, *b*_*c*_*f*. The network was trained to cross a threshold in anticipation of the time of the airpuff, however, no explicit form of the target function was imposed on the network. The target function was only defined at brief time points (10 ms) on three occasions, viz., at the mean of the distribution on the threshold, at the start of the trial at zero, and at the end of the trial at zero again. Despite the varied shapes of the training output solution by various networks, it was observed that expert networks followed the statistics of the stimulus for both paired and test trials. However, the statistics of the output of the switch networks were not different for the distribution conditions for paired and test trials, although their unit activities exhibited significant heterogeneity that matched that observed in neural responses. Network training was carried out in a manner similar to previous work^14,54^.

### Interval reproduction network datasets

In the third task, participants measured a sample interval and reproduced it immediately after measurement, similar to previous work^75^. On each trial, these intervals were sampled from a uniform prior distribution of time intervals (Prior). Previous work^55,75,76^ demonstrated that monkey and human timing responses during this task were biased towards the mean of this prior distribution in a manner predicted by Bayesian estimators (such as Bayes Least Squares or BLS). In the absence of prior knowledge, one would expect no bias towards the mean of the prior but an increase in variance, mimicking scalar variability in timing, as predicted by non-Bayesian models like Maximum Likelihood estimation (MLE)^57^. When neural activity was recorded in the macaque dorsomedial frontal cortex during this task, population dynamics assumed a semicircular trajectory, which when read out linearly gave rise to states that were compressed around the mean of the prior interval. These compressed neural states could account for both the Bayesian model predictions as well as the monkey’s responses, which were also consistent with Bayesian models. This leads to the hypothesis that the curvature of the semicircular trajectory was encoding prior knowledge, leading to compressed states when read out linearly. However, if the trajectory is unfolded this leads to the converse hypothesis that if instead the state estimates were read along the manifold geodesic, the outcomes would represent responses incurred in the absence or after the erasure of prior knowledge, consistent with Maximum Likelihood estimation (MLE).

We use the RNN to recreate this task and the underlying neural dynamics during the measurement epoch leading to the construction of a three-dimensional manifold, where the third dimension is orthogonal to the task-related manifold. Based on earlier work, it was expected that linearly decoding the curved manifold (by projecting onto its diameter) in its ambient dimension of three would yield readouts consistent with Bayesian models. On the other hand, embedding this manifold into two dimensions using manifold learning techniques and decoding along geodesic distance was expected to yield readouts consistent with MLE.

To achieve this, RNNs were trained using a two-dimensional time-varying input *u*(*t*) that consisted of a constant drive whose offset on each trial was sampled from a Gaussian distribution (𝒩0.5,0.1), a brief pulse input (20 ms) that demarcated the Start of the trial, after which a time interval was sampled from a discrete uniform distribution (529-1059 ms, comprising ten intervals), which was followed by another 20 ms pulse to mark the end of the time interval. The network was trained to cross an arbitrary threshold (*z*(*t*)=1) at the time of the sampled interval. During training, duration-dependent noise (Weber fraction dependent = 0.2) was introduced into the time of the target. Training under such scalar noise conditions induces the behavioral output to follow a compression towards the mean of the prior, which is the statistically optimal solution for such conditions.

## Footnotes

1 Note that RATS is not optimised on global distortion, which is an independent measure to evaluate the different manifold learning methods.

## Notes

### Competing Interest Statement

The authors have declared no competing interest.

### Summary of Updates

Improvements to method and demonstration on a wide range on complex neuroscience datasets. Also proposed, additional data pre-processing tools to make manifold discovery more accessible to the community.

