## Supplementary Information for "Unsupervised manifold learning using low-distortion alignment of tangent spaces"

### Supplementary figures

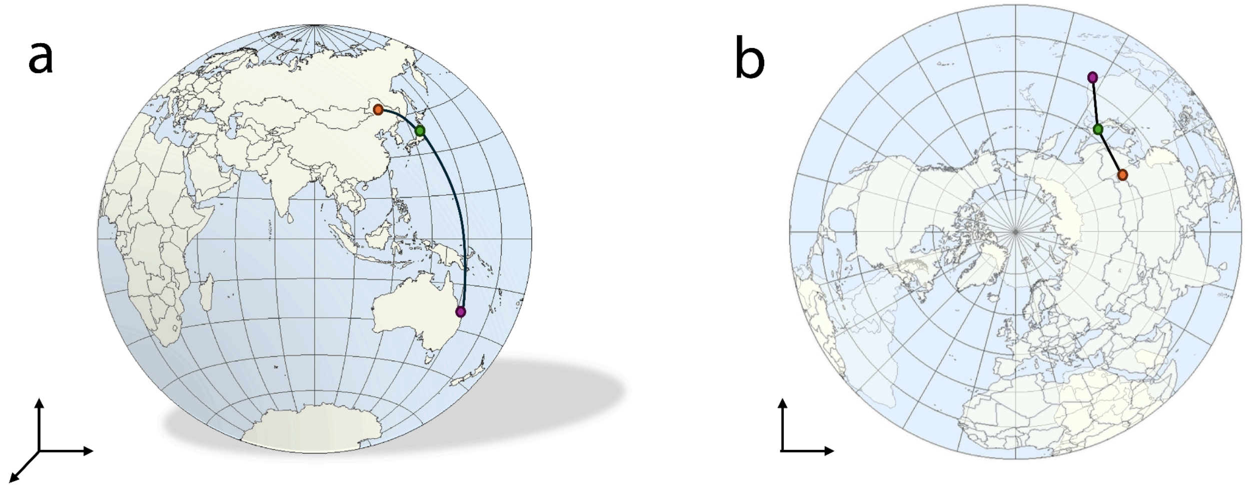


Supplementary figure 1: Illustration of distance distortion *a) The surface of the Earth on a 3D sphere. China (red) is close to Japan (green) and Australia (purple) is far. b) Rather than tearing the sphere to embed in two dimensions as with an atlas, a collapsed embedding overlays the north hemisphere on top of the south hemisphere in two dimensions. In this embedding Australia is as close to Japan as China is, indicating high distance distortion in the embedding.*

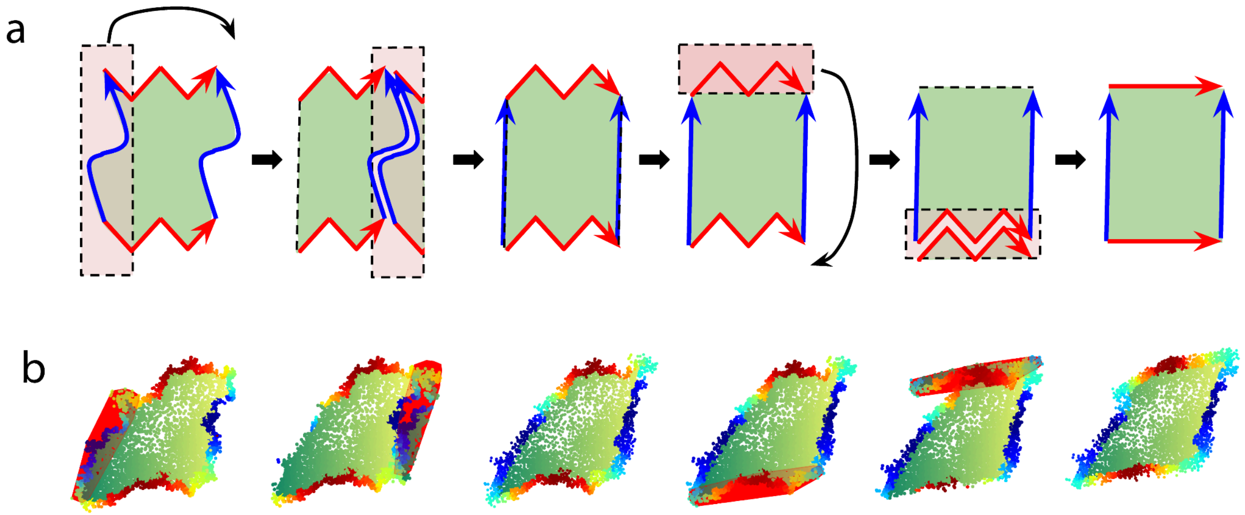


Supplementary figure 2: *Extensions to Gluing diagram. a) The schematic illustrates the process of transitioning between two gluing diagrams through a cut-and-paste technique. A segment of the embedding is sliced and subsequently glued along the arrows of matching colors. By pasting the cut segment along the same colored arrow, the topology of the embedding is maintained by appropriately updating the gluing diagram. The gluing diagram is recalculated and the process can be repeated. b) Using a GUI, we select a portion of the embedding on the left to*

*cut and reposition it closer to the area on the right where it is to be glued. The optimal alignment of the local views is recalculated using Riemannian gradient descent along with an update of the gluing instructions. The process is then repeated for a second time.*

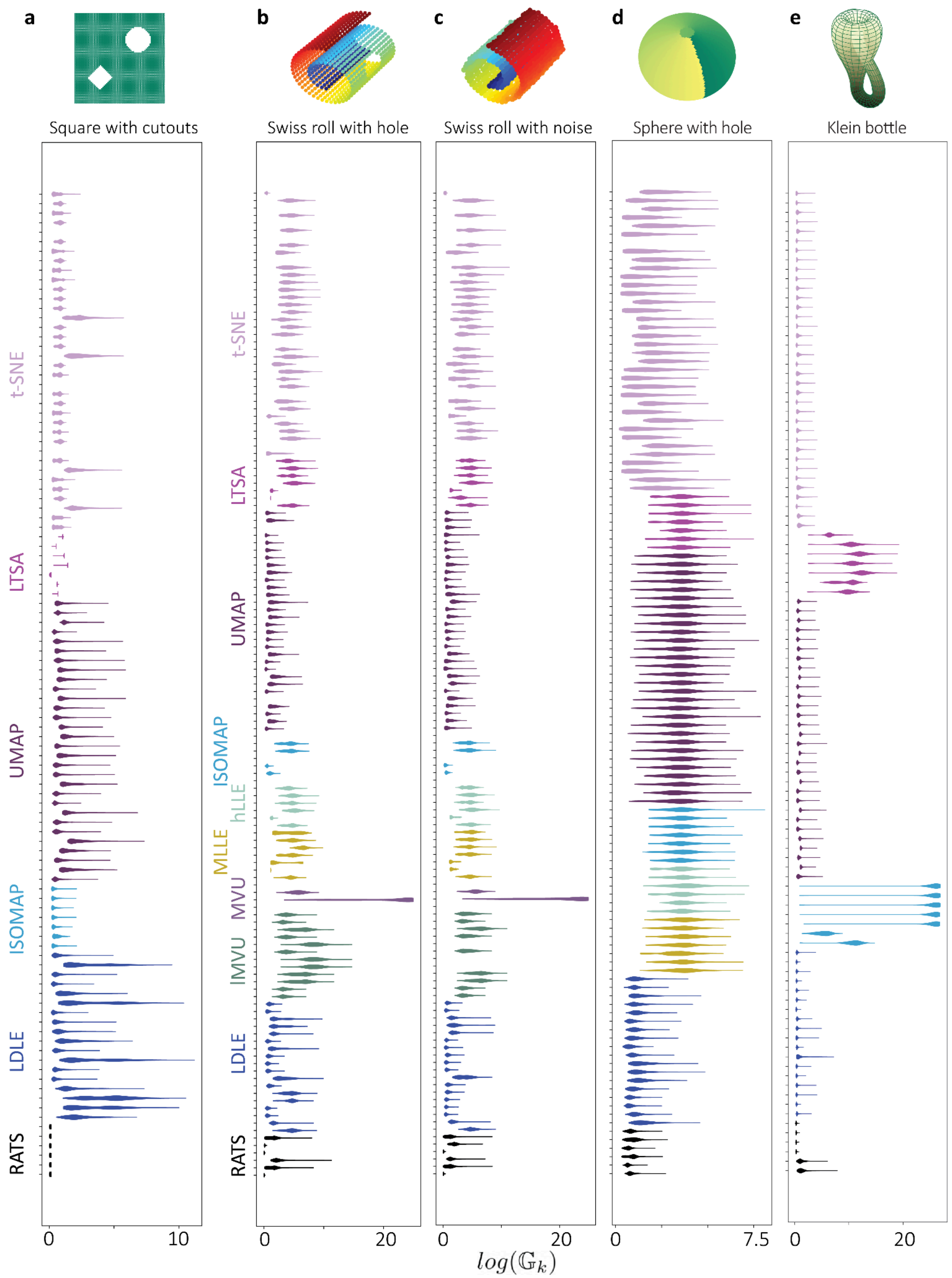

a
Square with cutouts
b
Swiss roll with hole
c
Swiss roll with noise
d
Sphere with hole
e
Klein bottle
t-SNE
LTSA
UMAP
ISOMAP
hLLE
MLLE
MVU
IMVU
LDLE
RATS
0
10
20
7.5
$log(\mathbb{G}_k)$

Supplementary figure 3: *Hyperparameter optimization for idealized datasets a) Square with cutouts b) Swiss roll with hole c) noisy swiss roll, d) sphere with a hole and e) Klein bottle. Violin plots representing the global distortion incurred by embeddings generated using various hyperparameter settings for each method. The absence of a violin plot for a hyperparameter setting indicates infinite global distortion, i.e.* $\mathbb{G}_k$ *is infinite for some* $k$.

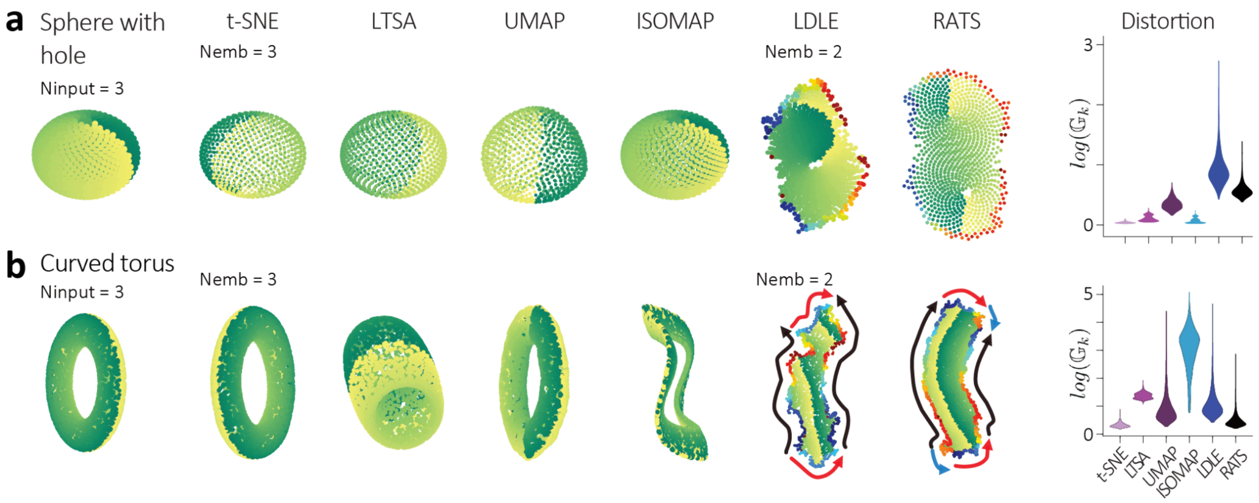


Supplementary Figure 4. Comparison of closed-manifolds with $N_{emb}$ disparity. a) Left: Sphere with a hole ($N_{input}$ = 3) is embedding in $N_{emb}$ = 3 using t-SNE, LTSA, UMAP, ISOMAP and in $N_{int}$ = 2 using LDLE and RATS. Right: Distortion plots. b) Left: Similar demonstration for a curved torus ($N_{input}$ = 3), embedded in $N_{emb}$ = 3 using t-SNE, LTSA, UMAP, ISOMAP and in $N_{int}$ = 2 using LDLE and RATS. Right: Distortion plots reveal that only t-SNE exhibits lower distortion whereas RATS, despite mandatory distortion by embedding in intrinsic dimension, shows the second-lowest distortion.

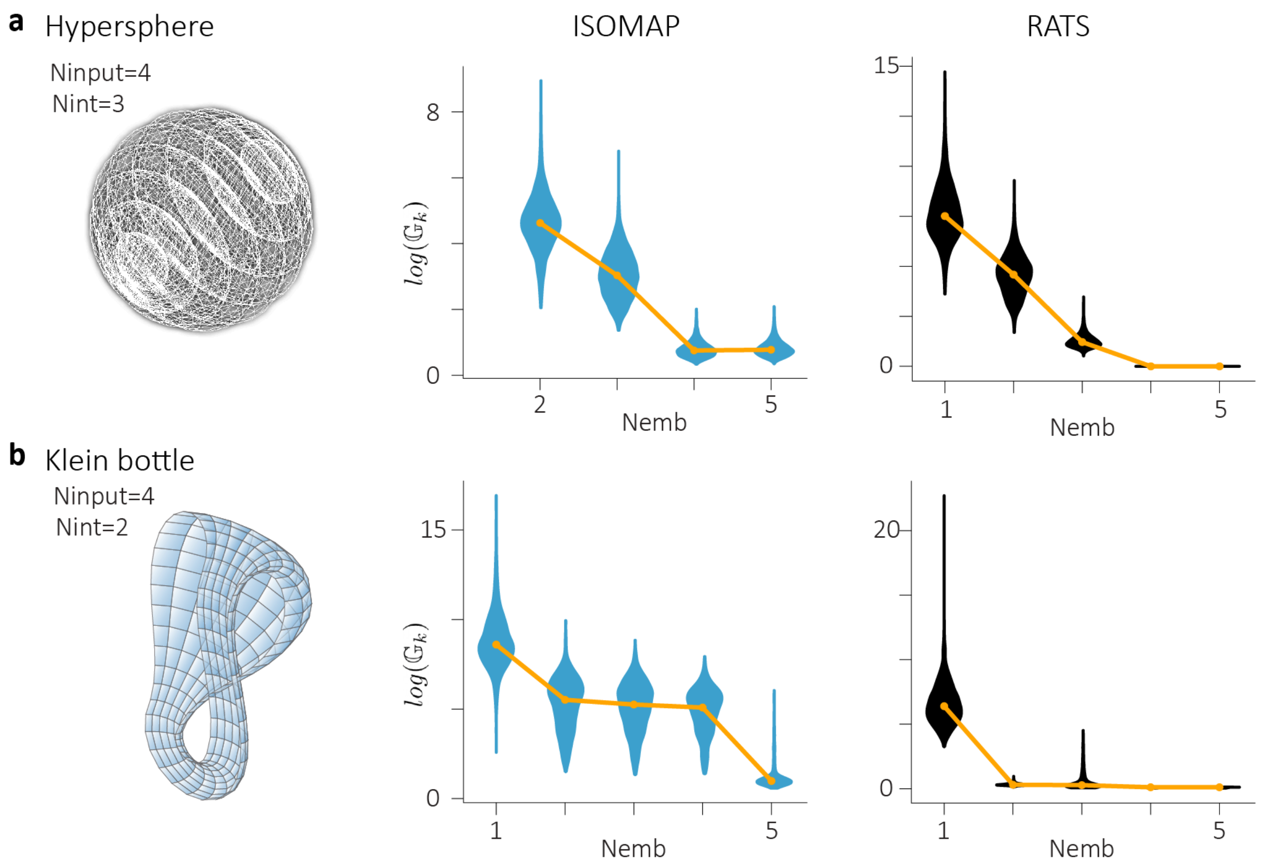


Supplementary figure 5: Using global distortion to estimate intrinsic dimensionality for Isomap (blue) and RATS (black) for a) 3-sphere (hypersphere) and b) Klein bottle.

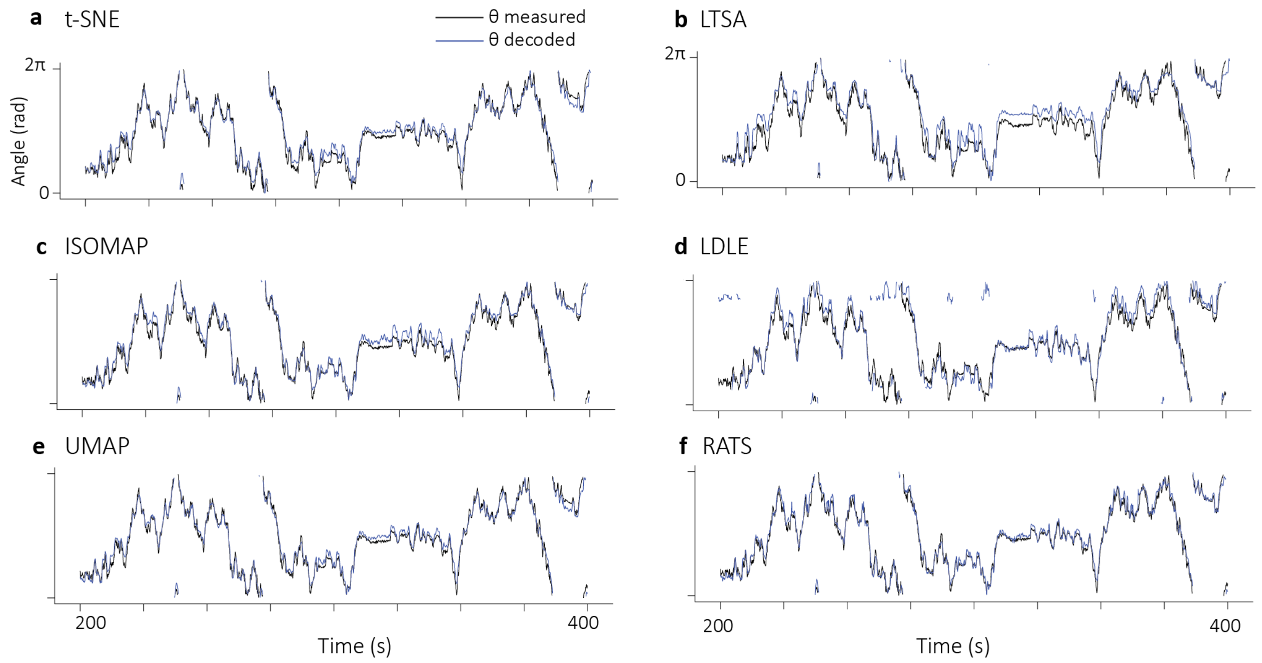

Supplementary figure 6: *Decoding of head direction. Measured head direction angle (black) and the decoded angle (blue) derived from embeddings generated by a) t-SNE, b) LTSA, c) ISOMAP, d) LDLE, e) UMAP, and f) RATS over time. Here t-SNE, LTSA, ISOMAP and UMAP are embedded in Nemb = 2 whereas LDLE and RATS are embedded in Nint = 1.*

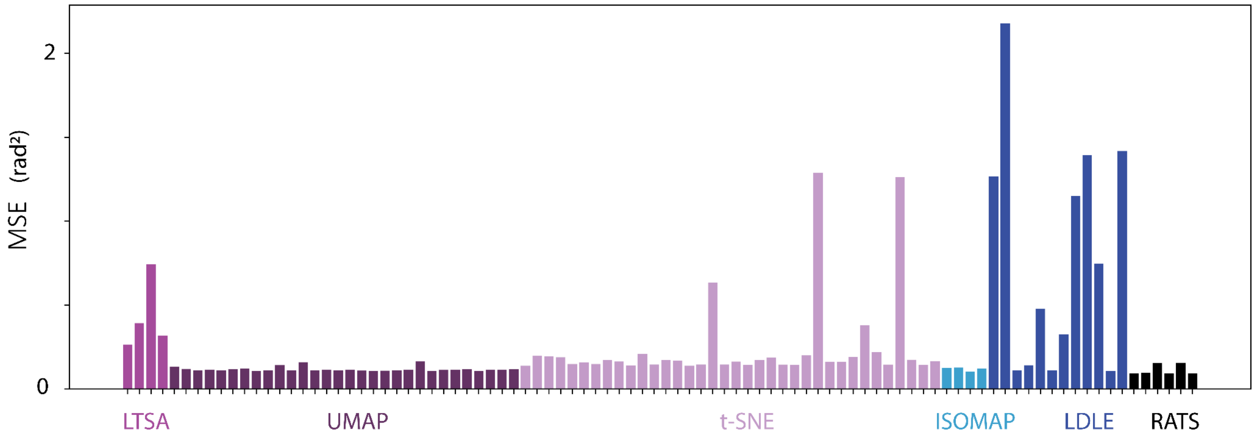


Supplementary figure 7: *Hyperparameter optimization for head direction dataset. Mean squared error (MSE) between the measured head direction angle and the decoded angle derived from embeddings generated using various hyperparameter settings for each method. Here, t-SNE, LTSA, ISOMAP and UMAP are embedded in Nemb = 2 whereas LDLE and RATS are embedded in Nint = 1.*

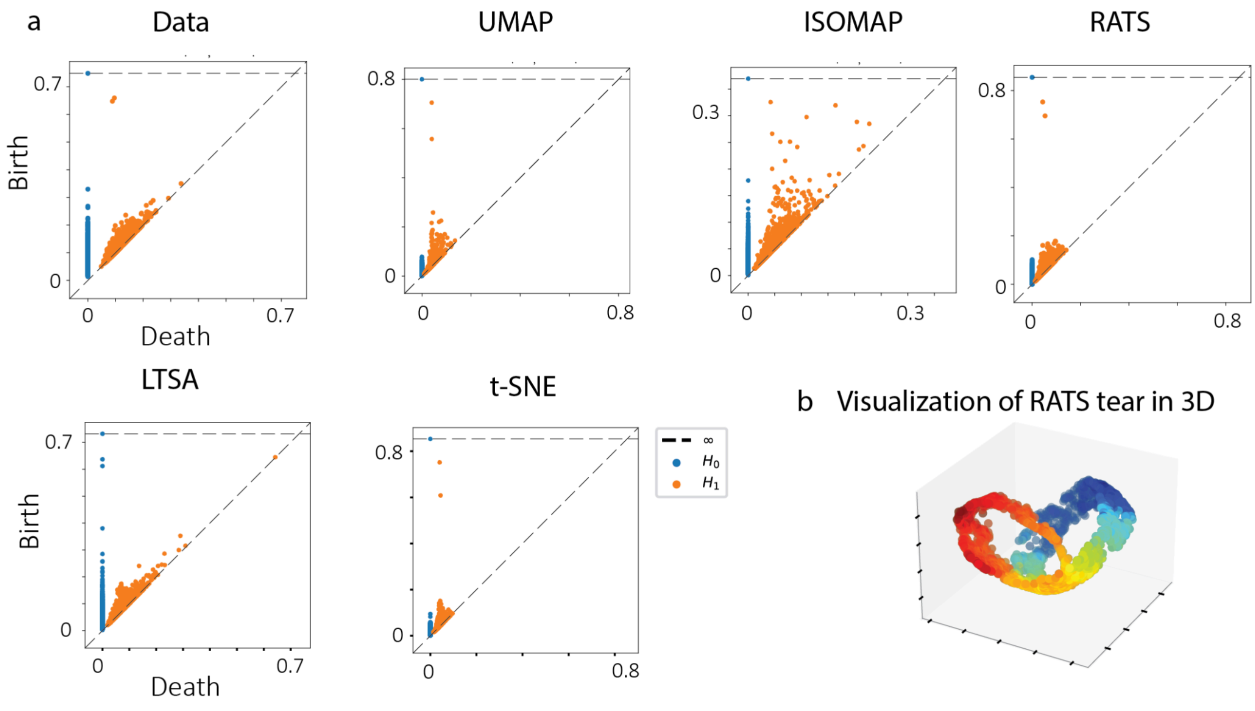

Supplementary figure 8: a) *Persistence diagrams for various embeddings provided in Figure 3F. b) Visualizing the points along the tear in the RATS embedding from Figure 3f on the UMAP embedding shows two circular loops, visually confirming the two H1 co-cycles identified in the persistent homology analysis.*

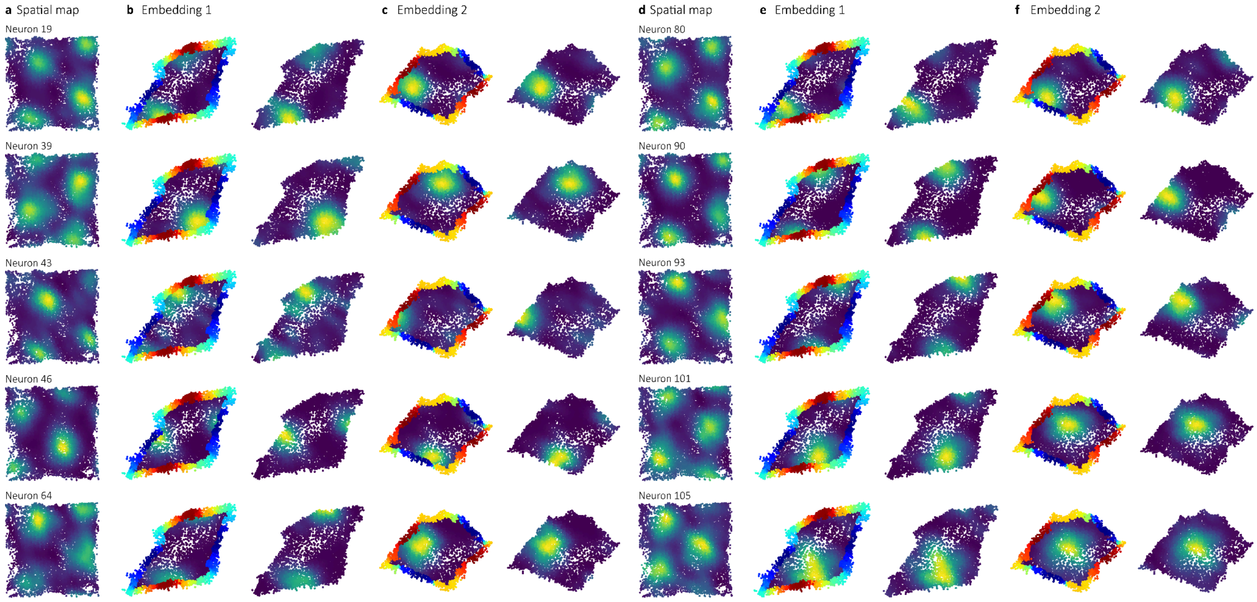


Supplementary figure 9: *Firing rate maps for selected grid cells across different RATS embeddings. The rate maps for various grid cells are shown in spatial coordinates (a,d) and on two distinct RATS embeddings where Embedding 1 (b,e) corresponds to figure 3F and embedding 2 (c,f) is obtained with a different hyperparameter set (still recovering a toroidal gluing diagram).*

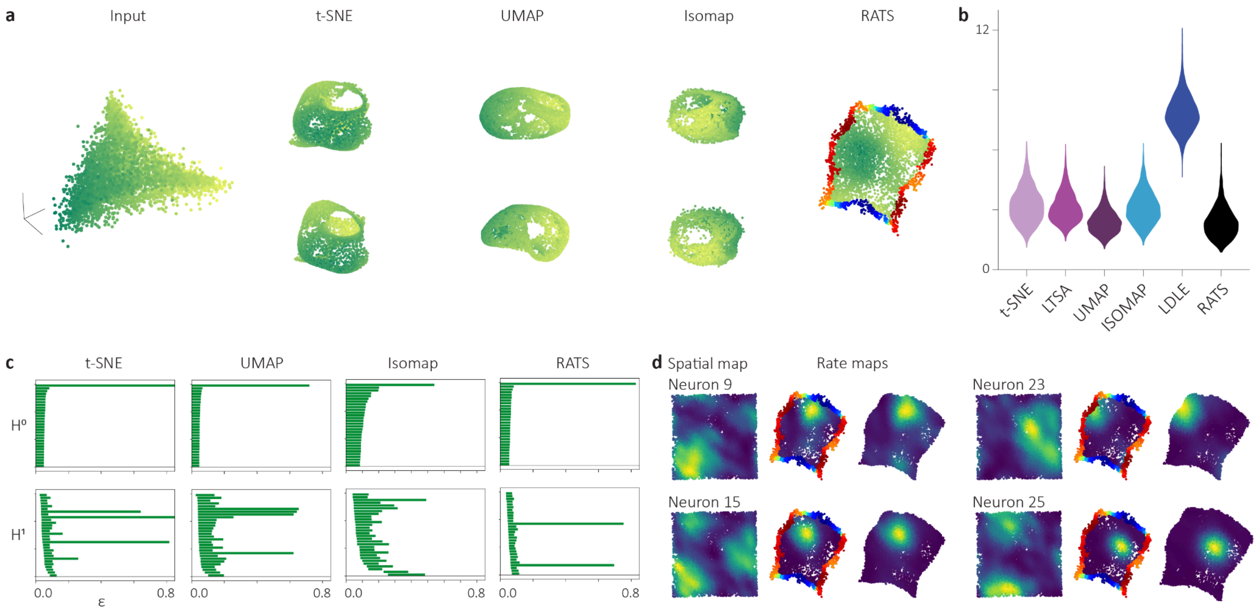


Supplementary figure 10: *Additional dataset example for* [13]. *a) The 145-dimensional population activity vectors recorded from MECII of a rat. Embeddings of the 6-dimensional PCA-reduced neural activity are obtained using t-SNE, LTSA, UMAP, and ISOMAP ($N_{emb}$ = 3) and using RATS ($N_{emb}$ =$N_{int}$ = 2). RATS provides gluing instructions by coloring the two sides of the 2D embedding. The derived gluing diagram depicts toroidal topology. b) Global distortion distributions (Statistics in Supplementary Table 2). c) The persistence barcodes for zeroth $H_0$ and first $H_1$ cohomology of the input data and the embeddings (see Methods). d) Visualization of the firing rate maps of four of the grid cells on the RATS embedding and in the spatial coordinates.*

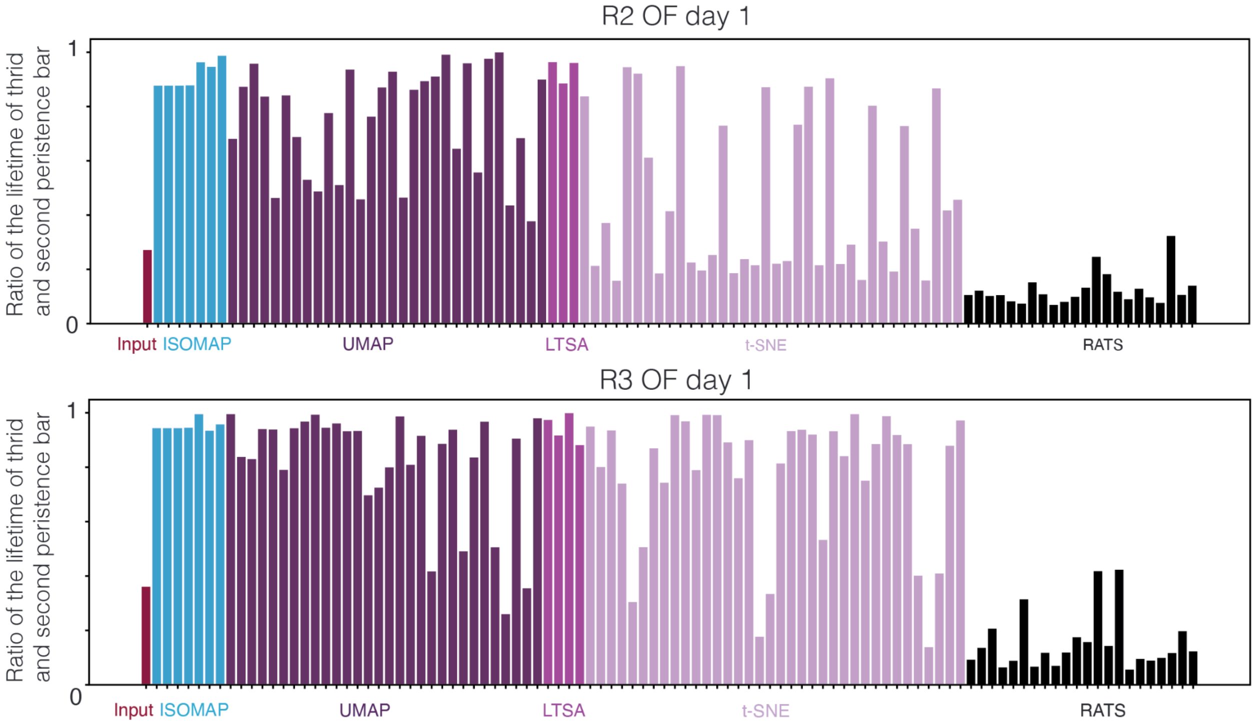


Supplementary Figure 11: *The ratio between the lifetimes of the third and second persistence bars in the* $H_1$ *barcodes, computed from both the input data (leftmost bar in the figure) and the embeddings of the grid cell dataset (Figure 3) as well as the additional grid cell dataset (Supplementary Figure 10), evaluated across different hyperparameter settings.*

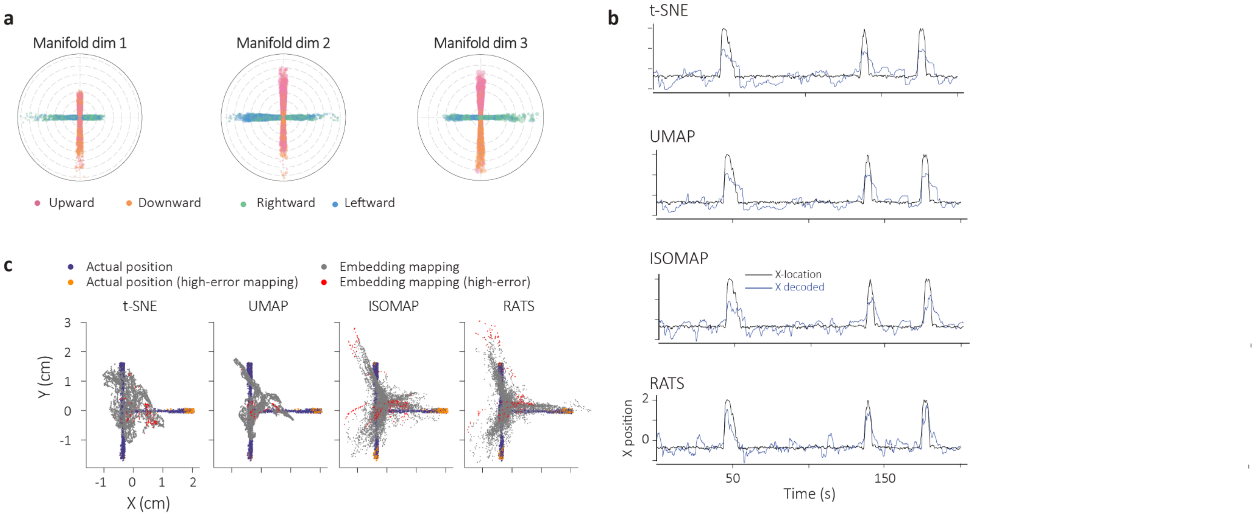


Supplementary figure 12: *Decoding from embedded data from ref*[50] *for the T-maze. a) Evidence that the intrinsic dimension of the T-maze embedding is 3. Decoding of speed in various directions in which the animal is moving along manifold dimension 1,2, and 3 reveal a clear demarcation of speed and direction related information in the*

*third dimension. b) Examples of decoded X-location from t-SNE, UMAP, ISOMAP and RATS. Black lines indicate actual X location and blue lines represent decoded X location. c) Mapping of different embeddings onto the 2D T Maze using t-SNE, UMAP, ISOMAP and RATS. Gray and red points indicate embedding samples corresponding to low and high mean squared error, respectively, along the first two manifold coordinates.*

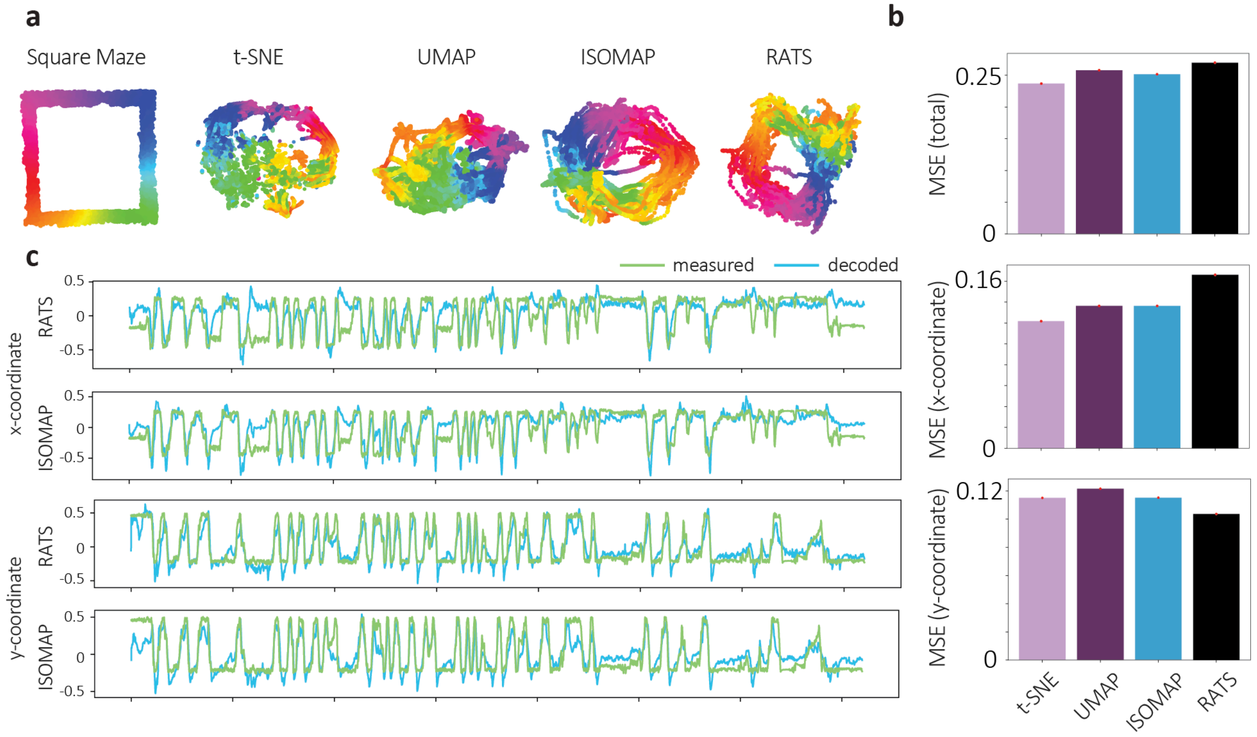


<u>Supplementary figure 13:</u> *Decoding from embedded data from ref[50] for square maze. a) Embeddings of 10-dimensional PCA reduced calcium imaging data (N = 115 units) using t-SNE, UMAP, ISOMAP and RATS. Colors in embeddings (right) correspond to locations on the square maze (left). b) mean squared error (total, x-coordinate only and y-coordinate only) for different methods when comparing actual and mapped locations from the embeddings. Statistics can be found in Supplementary table 2. c) The actual and the mapped locations obtained from RATS and ISOMAP embeddings.*

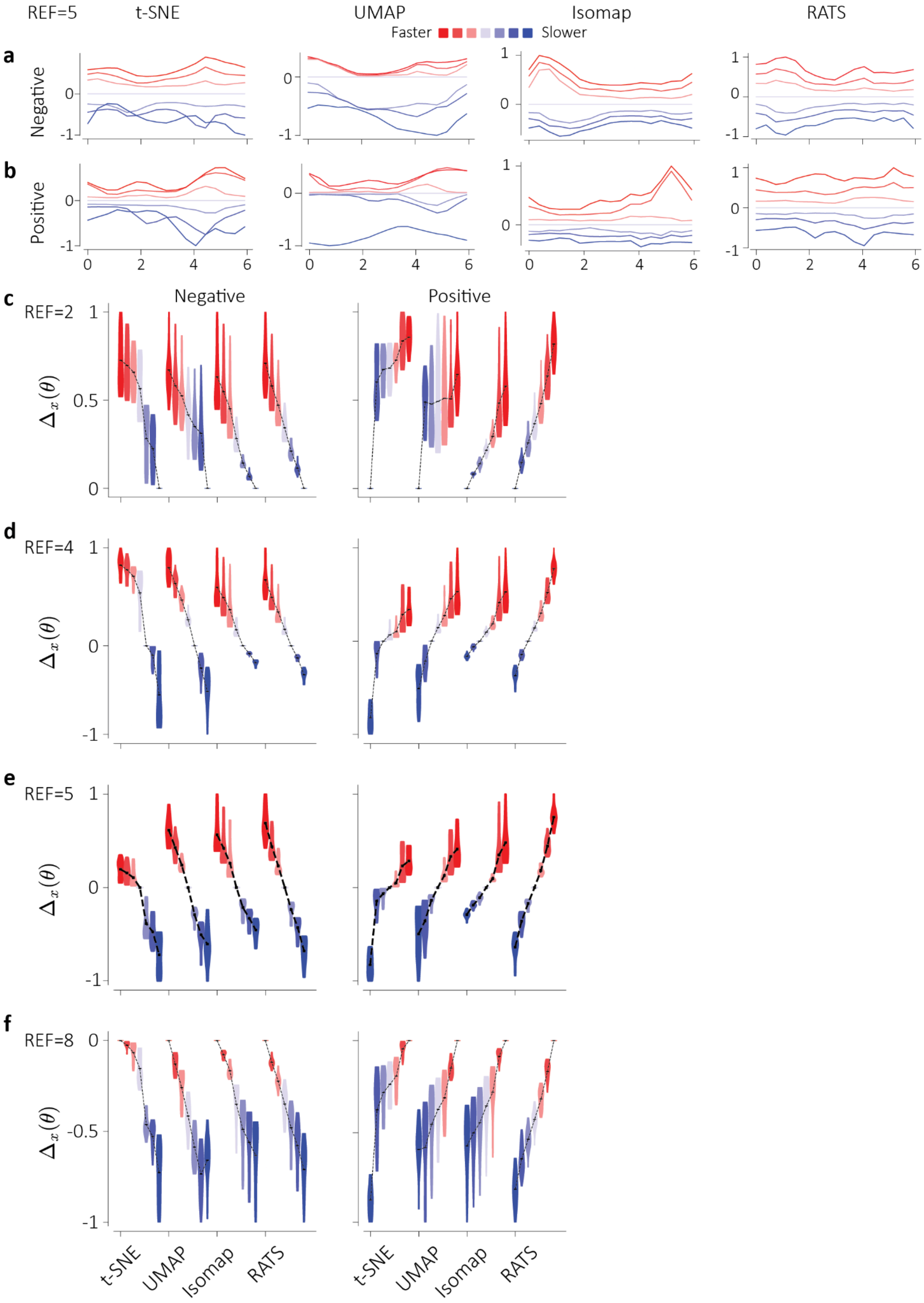

REF=5
t-SNE
UMAP
Isomap
RATS
Faster
Slower
a
Negative
b
Positive
c
REF=2
Negative
Positive
$\Delta_x(\theta)$
d
REF=4
e
REF=5
f
REF=8
t-SNE
UMAP
Isomap
RATS

Supplementary figure 14: *a) Angle-dependent distance of the reference trajectory (speed bin 4) from each of the other speeds for negative cycling direction for UMAP, t-SNE, Isomap and RATS embeddings. b) same as a) but for positive direction. c-f): Distribution of angle-dependent distances between reference trajectory at given speed bin and each of the other speeds for negative (left) and positive (right) cycling directions.*

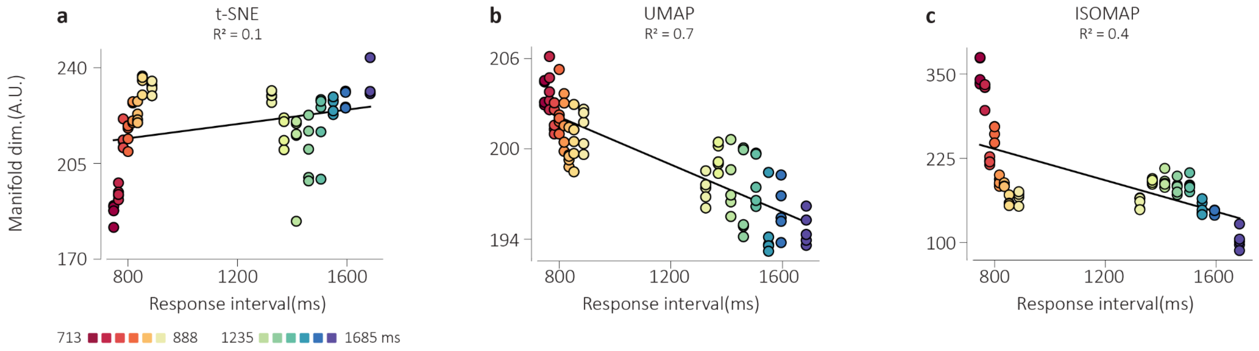


Supplementary Figure 15: *Decoding of thalamic single unit data for response interval for A) t-SNE, B) UMAP and C) Isomap. Red colors indicate intervals in the range 713-888 ms, blue colors indicate intervals in the range 1235-1685 ms. Black lines indicate linear fits with coefficient of determination reported. Statistics can be found in Supplementary table 5.n*

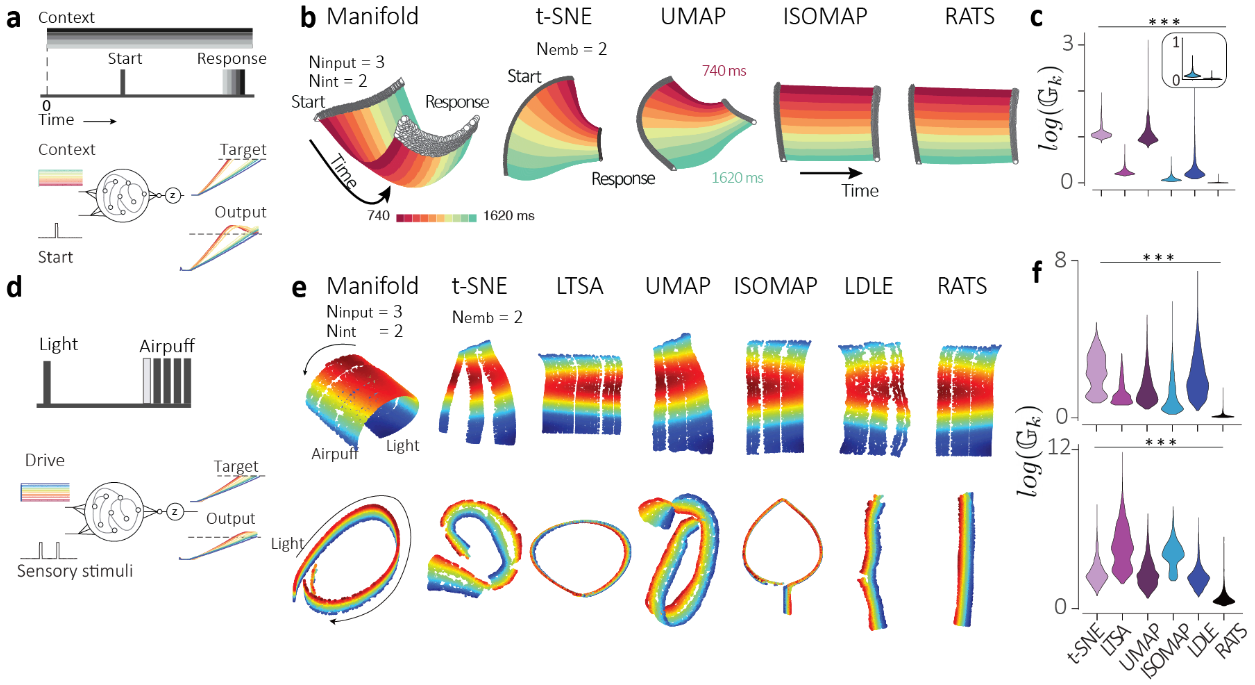


Supplementary Figure 16: *Embedding irregular manifolds generated through deep networks. a) Based on ref[54], RNNs were trained to associate context cues with intervals by ramping to a threshold at different times. b) Left: Nonlinear manifold obtained from the network dynamics (left) embedded using t-SNE, UMAP, ISOMAP and RATS.*

*Colors represent context-dependent time interval associations (red to green: 740 - 1620 ms). Right: Global distortion distributions for embeddings of the nonlinear motor timing manifold using t-SNE, LTSA, UMAP, ISOMAP, LDLE and RATS. Non-parametric analysis of variance using Kruskal-Wallis . c) Left: In ref [33], neural population activity speeds for different contexts correlated with the produced time interval (red to violet: 740 - 1620 ms). Right: Nonlinear manifolds generated by dynamics of trained RNNs recapitulate this principle, where despite variation in density of the manifold for short and long durations, decoding of the manifold yields a high correlation with produced time intervals. d)  Manifolds generated by RNNs trained on an associative learning task between two sensory signals (light and airpuff)[52]. RNNs are provided a continuous driving signal that represents fluctuations in initial condition of the dynamics. e) Resulting nonlinear manifolds ($N_{input}$ = 3) containing missing data and containing a forked tail are embedded using t-SNE, LTSA, UMAP, ISOMAP, LDLE and RATS. f) Global distortion distributions for each embedding for the manifolds.*

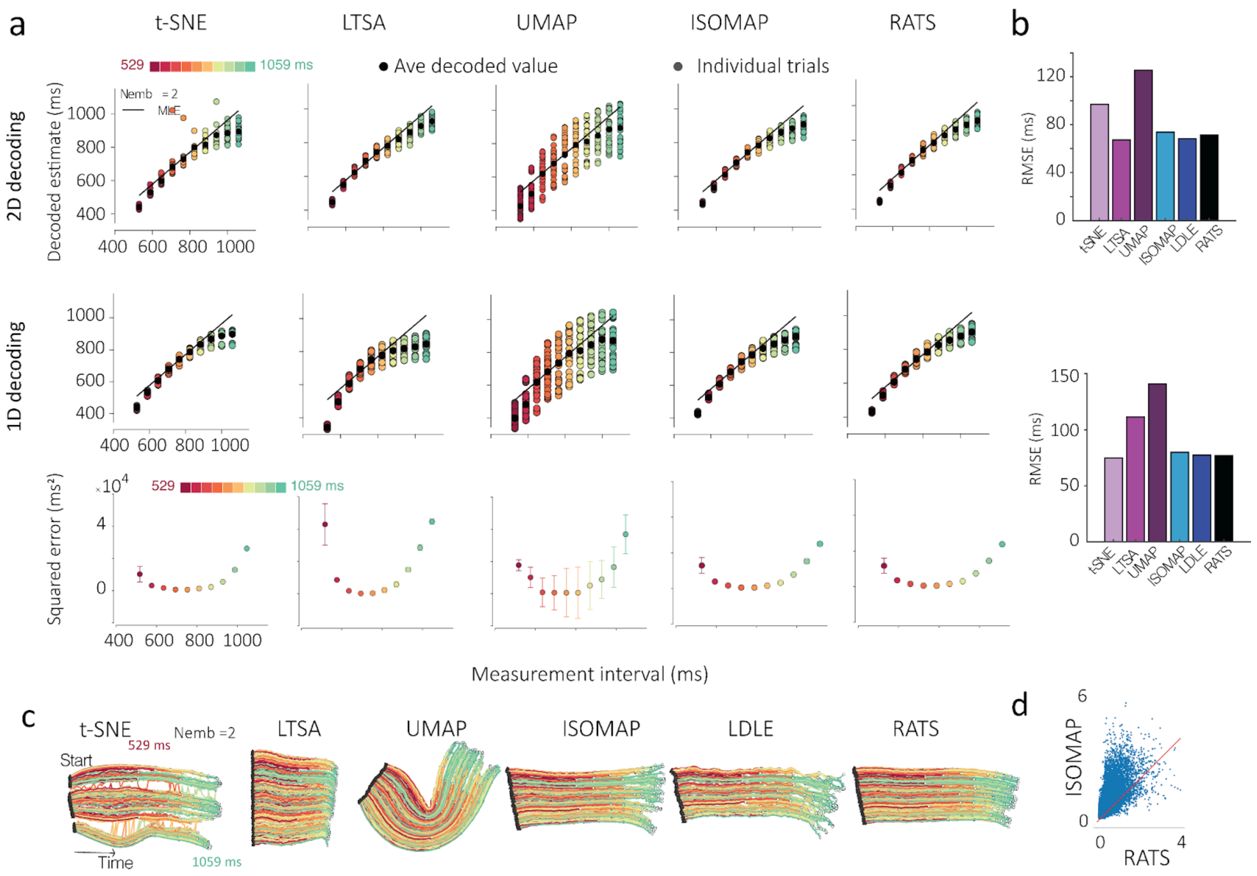


<u>Supplementary figure 17: Time interval reproduction task.</u> a) one and two-dimensional decoding of time interval estimates along the embedding geodesics and comparison with measurement time provided to the network on a trial-by-trial basis. Decoding is performed for t-SNE, LTSA, UMAP, ISOMAP, and LDLE. Colors indicate uniformly sampled intervals from 529 - 1059 ms (red to green). Colored circles represent decoding on individual trials, black circles represent averages. Error bars indicate standard error of the mean. Lowest row illustrates per condition

squared error in 1D decoding estimates for various techniques. Error bars indicate standard deviation. b) Root mean squared error (RMSE) comparisons for each technique's decoded estimates (2D above, 1D below) with the measured estimates for all techniques. c) In addition to embeddings shown in main figure 2c, embeddings ($N_{emb}$ = 2) of the input ($N_{input}$ = 3) is shown here for all techniques. Triangles indicate start points and circles indicate termination of the interval. Color scheme is the same as in a). d) Log of point-wise global distortion due to embedding with ISOMAP vs. RATS.

### Supplementary tables

| Dataset | N | DOF = #methods-1 | H Statistic | P value |
|---|---|---|---|---|
| Barbell | 4929 | 5 | 15815.19 | P < 0.001 |
| Square with two holes | 4543 | 5 | 19600.07 | P < 0.001 |
| Swissroll with hole | 960 | 9 | 8761.16 | P < 0.001 |
| Noisy Swiss roll | 990 | 9 | 8923.54 | P < 0.001 |
| Sphere with hole | 993 | 9 | 4620.36 | P < 0.001 |
| Curved torus 2d embeddings | 5000 | 5 | 24005.40 | P < 0.001 |
| Klein bottle | 4900 | 5 | 24652.19 | P < 0.001 |
| Sphere with hole 3d embeddings | 960 | 9 | 5183.35 | P < 0.001 |
| Curved torus 3d embeddings | 5000 | 5 | 24368.22 | P < 0.001 |

Supplementary table 1: Kruskal-Wallis statistics for idealized datasets in Figure 2.

| Dataset | N | DOF=#methods-1 | Statistic | P value |
|---|---|---|---|---|
| Head direction cells population | 5000 | 5 | H = 49.36 | P < 0.001 |
| Grid cells population (R2) | 4800 | 5 | H = 8058.99 | P < 0.001 |
| Grid cells population (R3) | 4800 | 5 | H = 10258.34 | P < 0.001 |
| Maze exploration (T-Maze) | 7333 | 3 | F = 5.2672 | P < 0.05 |
| Motor cycling | 7051 | 3 | F=526.58 | P < 0.001 |

Supplementary table 2: Kruskal-Wallis statistics for Figure 3 and Figure 4 datasets. Degrees of freedom=5 (#methods-1).

| Linear regression | t-SNE | UMAP | ISOMAP | RATS |
|---|---|---|---|---|
| R square | 0.10 | 0.68 | 0.40 | 0.69 |
| P value | 0.003 | <0.001 | <00.1 | <0.001 |
| R squared | 0.751 | 0.684 | 0.651 | 0.673 |
| P value | <0.001 | <0.001 | <0.001 | <0.001 |

Supplementary table 3: Co-efficient of determination statistics for various methods and datasets for motor timing and trace conditioning.

| Dataset | N | DOF=#methods-1 | H Statistic | P value |
|---|---|---|---|---|
| Motor timing | 10431 | 5 | 56204.16 | P < 0.001 |
| Trace conditioning 1 | 6375 | 5 | 21667.80 | P < 0.001 |
| Trace conditioning 2 | 6650 | 5 | 25361.11 | P < 0.001 |
| Interval reproduction | 12796 | 5 | 39923.16 | P < 0.001 |

Supplementary table 4: Kruskal-Wallis statistics for datasets in Supplementary figure 16-17. Degrees of freedom=5 (#methods-1).

| Wilcoxon test | RATS vs t-SNE | RATS vs LTSA | RATS vs UMAP | RATS vs ISOMAP | RATS vs LDLE |
|---|---|---|---|---|---|
| W statistic | 0 | 0 | 0 | 64673 | 6 |
| P value | P < 0.001 | P < 0.001 | P < 0.001 | P < 0.001 | P < 0.001 |

Supplementary table 5: Two-sided Wilcoxon signed-rank test statistics for the motor timing task (Figure 4).

| Wilcoxon test | t-SNE vs UMAP | ISOMAP vs t-SNE | ISOMAP vs UMAP | RATS vs t-SNE | RATS vs UMAP | RATS vs ISOMAP |
|---|---|---|---|---|---|---|
| W statistic | 8 | 63 | 0 | 140 | 0 | 339 |
| P value | P < 0.001 | P < 0.001 | P < 0.001 | P < 0.001 | P < 0.001 | P < 0.001 |

Supplementary table 6: Two-sided Wilcoxon signed-rank test statistics for the Trace conditioning task RMSE
